# Temperature and precipitation interactively shape the plant microbiome by regulating the start of the growing season

**DOI:** 10.1101/2025.01.23.634453

**Authors:** Dina in ‘t Zandt, Anna Florianová, Mária Šurinová, Michiel H. in ‘t Zandt, Kari Klanderud, Vigdis Vandvik, Zuzana Münzbergová

## Abstract

Climate change is altering associations between plants and soil microbiota, threatening ecosystem functioning and stability. Predicting these effects requires understanding how concomitant changes in temperature and precipitation influence plant-soil microbiota associations. We identify the pathways via which temperature and precipitation shape prokaryote and fungal rhizosphere and root-associated communities of the perennial grass *Festuca rubra* in cold-climate ecosystems. We found that interactive effects of temperature and precipitation are key in shaping plant-soil microbiota associations, with the start of the growing season as a critical mediating factor. Specifically, the start of the growing season is advanced by increasing temperature, but delayed by increasing precipitation. This interactive pathway particularly shaped rhizosphere organic matter degrading microbiota, and root-associated plant pathogen and beneficial microbiota. We conclude that understanding local temperature, precipitation, and seasonal changes is crucial to accurately predict how the unique plant-microbiota interactions shaping cold-climate ecosystems are evolving with the ongoing change in climate.

## Introduction

The ongoing changes in climate are altering interactions between plants and soil microbiota. Established interactions are being disrupted, while novel interactions emerge^1^. These shifts in plant-soil microbiota interactions pose a threat to the functioning and stability of ecosystems, as these interactions are a critical driver of numerous plant community processes, including succession, coexistence, invasion, range expansion, diversity, and stability^2–9^. Climate change impacts are particularly concerning in cold-climate ecosystems, where warming is occurring faster than the global average, and biological processes are often limited by low temperatures^10^. As these ecosystems warm, the conditions that plant-soil microbiota interactions are adapted to are lifted, likely resulting in profound consequences for overall ecosystem functioning.

At the same time, precipitation patterns in cold-climate regions are also changing. These concurrent shifts in precipitation and temperature are likely to have complex effects on plant-soil-microbiota interactions^10^. However, studies often focus on a single climate factor at a time, leaving a significant gap in our understanding of how simultaneous changes in temperature and precipitation impact these interactions. This knowledge gap limits our ability to predict the broader consequences of climate change on cold-climate ecosystem functioning and stability. In this study, we fill this knowledge gap by identifying how concomitant changes in temperature and precipitation shape plant-soil microbiota interactions in cold regions.

Interactions between plants and soil microbiota occur in the rhizosphere, the thin layer of soil surrounding plant roots, as well as on the root surface (rhizoplane) and inside the root (endosphere)^11,12^. These interactions are sensitive to climate change, as even minor climatic changes trigger rapid physiological responses in plants and strong alterations in soil microbial community composition^1,13^. While the effects of warming on plant-soil microbiota interactions in cold regions are relatively well understood, the role of precipitation and its interaction with warming has received considerably less attention. This is largely because studies typically focus on the presumed limiting factor of a system, such as temperature in cold climates and precipitation in dry regions. Additionally, precipitation is inherently more complex than temperature, because it can fall as either rain or snow. This means that the consequences of changing precipitation patterns depend on when in the season precipitation changes, and thus whether snowfall or rainfall is affect, but also whether temperature changes concomitantly, potentially altering the form of precipitation. Not only do temperature and precipitation jointly determine soil moisture, their interactive effects also determine snow depth, the timing of snowmelt, and thus the onset of the growing season in cold-climate ecosystems. Given this complexity, understanding how simultaneous changes in both temperature and precipitation affect plant-soil microbiota interactions is essential for predicting impacts of climate change on cold-climate ecosystems.

In cold-climate ecosystems, warming raises the metabolic activity of both plants and soil microbiota. In plants, warming increases photosynthetic activity, leading to greater carbon (C) allocation belowground through increased root growth and root exudation^14–18^. This increase in belowground C allocation affects plant interactions with soil microbiota by, for example, enhancing rhizosphere priming: the promotion of rhizosphere microbiota performing mineralisation to increase nitrogen (N) availability^17,19,20^. In soil microbial communities, a raise in nutrient availability favours fast-growing microbiota over slow-growing microbiota^21^. This shift typically results in microbial communities becoming increasingly dominated by faster-growing prokaryotes compared to the generally slower-growing fungi^22^. A dominance of fast-growing microbiota is associated with accelerated soil C and N cycling, resulting from increased soil organic matter turnover with warming^14,23,24^. Consequently, warming of cold-climate ecosystems impacts both plant and microbial performance individually.

However, with changing precipitation, warming effects on plant-soil microbiota interactions are also likely to change. With the overall low metabolic activity and low evapotranspiration in cold conditions, precipitation increases soil moisture more significantly in cold than in warm conditions^24^. Consequently, in warm conditions, an increase in precipitation may fuel the higher metabolic rates of both plant and microbial communities, amplifying the effects of warming on plant, soil and microbial processes. In contrast, in cold conditions, increased precipitation may lead to more frequent anoxic soil conditions. Anoxic conditions severely impact both plant and microbial performance. For example, soil anoxia reduces soil microbial respiration and decomposition rates, and favours soil denitrification over nitrification pathways^25,26^. In plants, soil anoxia reduces CO_2_ assimilation and nutrient uptake, while also triggering anaerobic root metabolism and changes in root traits, such as root tissue density^27–29^.

On a seasonal scale, changes in precipitation – specifically snowfall – are likely to affect how warming shapes plant-soil microbiota associations by influencing the start of the growing season. While an increase in temperature accelerates snowmelt and advances the start of the growing season, an increase in precipitation increases snow depth and delays the start of the growing season^30^. Snow removal and snow addition experiments demonstrate the key role that advances and delays in the start of the growing season play in shaping soil biogeochemical and soil microbial communities^31–33^. In cold-climate ecosystems, temperature and precipitation changes are thus likely to interactively reshape plant-soil microbiota associations. However, we currently lack a clear understanding of the mechanisms by which these factors jointly influence these associations.

Beyond direct effects of warming, precipitation and their interactions on plant-soil microbial associations, climate change in cold-climate ecosystems is also likely to extensively alter plant-soil microbiota associations through indirect pathways. These indirect pathways may include shifts in plant community composition, soil biogeochemical processes, and changes in the neighbouring plant species with which host plants interact. For instance, warming of cold ecosystems increases plant community productivity, favouring fast-growing plant species and leading to the exclusion of slow-growing species, thereby decreasing plant community diversity^34,35^. Such changes in plant community composition and diversity may, in turn, influence plant-soil microbiota associations by altering soil biogeochemical processes and modifying the neighbouring species with which a host plant interacts^36^. We currently lack insight in the extent to which direct versus indirect climate-driven pathways mediate the effects of simultaneous changes in temperature and precipitation on plant-soil microbiota associations.

We test how prokaryote and fungal rhizosphere and root-associated (rhizoplane and endosphere) communities of *Festuca rubra* in cold-climate ecosystems are affected by temperature, precipitation and their interaction. *F. rubra* is a widely distributed perennial grass species growing in multiple climatic zones. This species therefore provides a unique opportunity of testing shifts in rhizosphere and root-associated microbiomes along large climatic gradients. We sampled *F. rubra* in the Vestland Climate Grid located in the fjords of southern Norway^35,37^. The grid comprises of 12 natural grassland locations differing in average summer temperatures (6.5, 8.5 and 10.5 °C) and average annual precipitation (600, 1200, 2000 and 2700 mm) in a factorial design^35,37^. We determined rhizosphere and root-associated prokaryote and fungal communities of eight *F. rubra* individuals from each location using 16S and ITS amplicon sequencing. Additionally, we measured soil abiotic properties and determined aboveground plant community composition. We ask (i) what the relative importance is of temperature, precipitation, and their interaction in shaping rhizosphere and root-associated prokaryote and fungal community composition, (ii) via which pathways these effects occur: directly as a response to changes in climate and seasonality, or indirectly, mediated by changes in plant community composition and/or soil abiotic properties, and (iii) which soil microbial groups are reshaped by these climate-driven pathways. Based on our findings, we discuss the effects of the ongoing concomitant changes in temperature and precipitation on plant-soil microbiota associations in cold-climate ecosystems, and the implications these shifts are likely having on ecosystem functioning and stability.

## Results

*Temperature, precipitation and their interaction shape the rhizosphere and root microbiome* Rhizosphere and root-associated prokaryote and fungal communities of *Festuca rubra* showed significant separation between the 12 locations (Fig 1A-D). Separation of rhizosphere and root-associated communities was significantly related to temperature, precipitation, and the interaction between temperature and precipitation (Fig 1A-D).

**Fig. 1.**
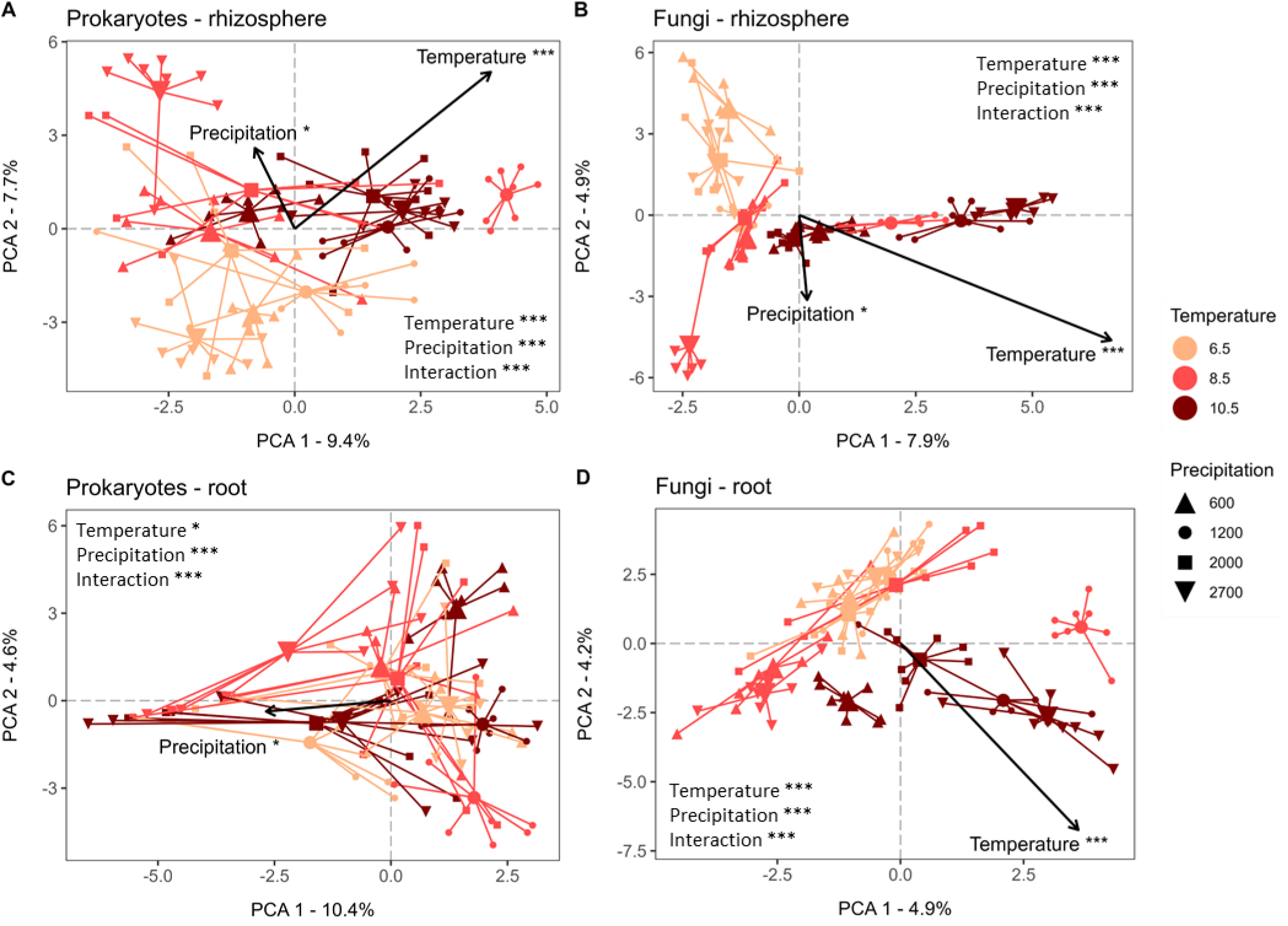
Principal Component Analysis (PCA) illustrating temperature and precipitation effects on plant-soil microbiota associations. (A) Rhizosphere prokaryotes, (B) rhizosphere fungi, (C) root-associated prokaryotes, and (D) root-associated fungi of *Festuca rubra*. Arrows indicate increasing temperature and precipitation. Colours indicate the temperature gradient (average summer temperature in degrees Celsius) and shapes the precipitation gradient (average annual precipitation in mm). Small shapes indicate the position of each *F. rubra* individual and large shapes indicate the centroids of each location (n = 8 rhizosphere and root samples per location with 12 locations in total). Results of PERMANOVA tests on significant separation of temperature, precipitation and their interaction are presented. Significance codes: *** = p < 0.001; * = 0.01 < p < 0.05.

Prokaryote communities were highly unique between the rhizosphere and root compartments. Only 17% of prokaryote taxa were observed in both the rhizosphere and root compartments (Fig S1AB). Within each compartment, however, a large core community of prokaryotes was present. Between 84 and 79% of prokaryote taxa were present at all three temperature levels, and between 77 and 63% at all four precipitation levels in the rhizosphere and root compartments, respectively (Fig S2AB, S3AB).

Fungal communities, on the other hand, showed an overlap of 85% in taxa occurrence between the rhizosphere and root compartments (Fig S1AB). Fungal taxa occurrence was more variable than for the prokaryote community with only 33 and 49% present at all three temperature levels, and 26 and 38% at all four precipitation levels for the rhizosphere and root compartments, respectively (Fig S2CD, S3CD).

### Microbial co-occurrence networks were affected by temperature, precipitation and their interaction

We created microbial co-occurrence networks for the rhizosphere and root compartments separately. In each network, we identified network clusters, which grouped similarly responding prokaryote and fungal taxa (Fig 2; Fig S4-S5). Importantly, both the rhizosphere and root co-occurrence networks were significantly more densely clustered than randomised networks, showing that a distinct organisational structure occurred in both rhizosphere and root-associated microbial networks (Fig S6).

**Fig. 2.**
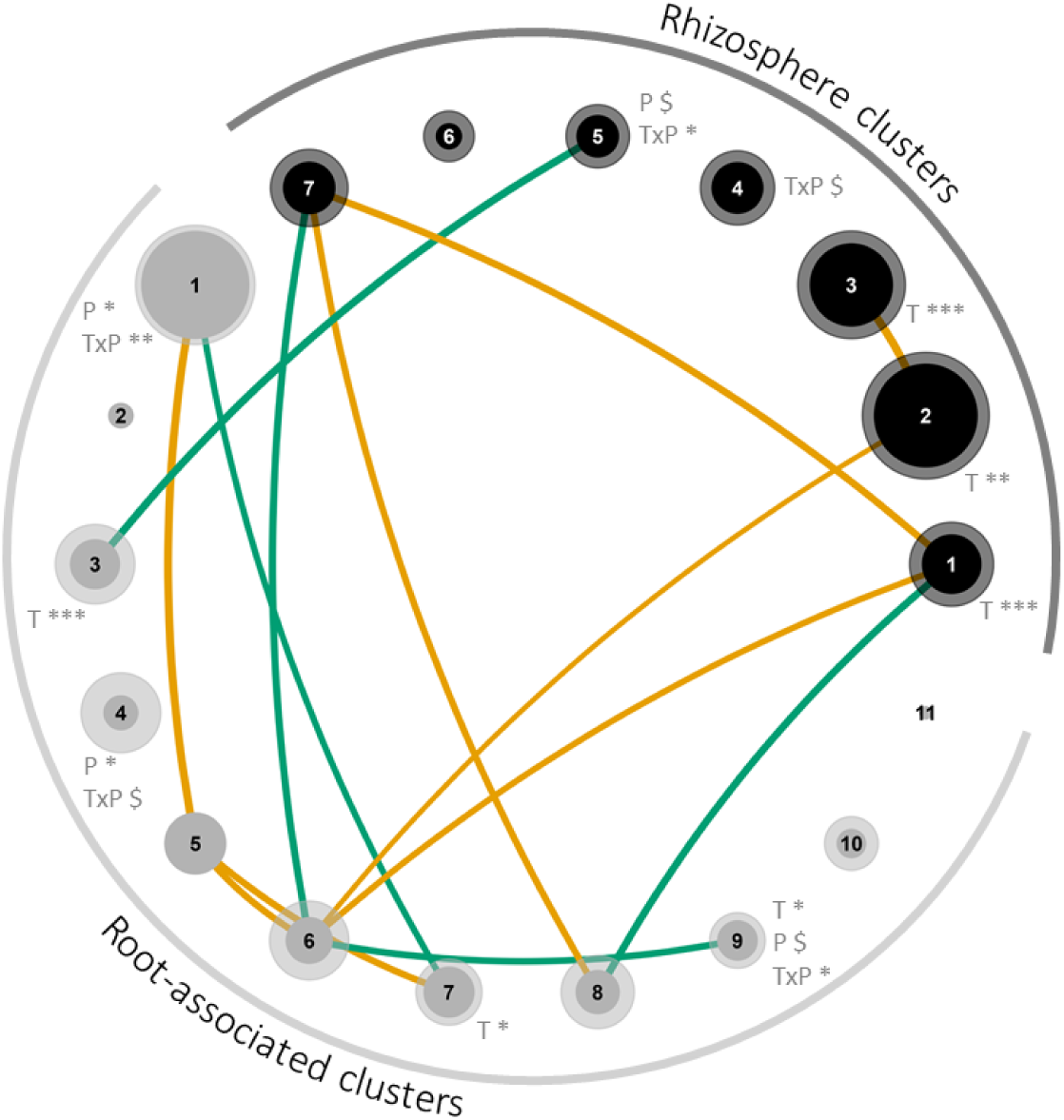
Co-occurrence network clusters of rhizosphere (dark grey) and root (light grey) microbial communities. Solid, central zones of each circle indicate the average relative abundance of prokaryote taxa in the cluster. Transparent, outer zones of each circle indicate the average relative abundance of fungal taxa in the cluster. Root-associated clusters 2, 5 and 11 contain prokaryote taxa only. Significant correlations between clusters are indicated by lines between clusters. Negative correlations are indicated in yellow, positive correlations in green. Width of lines indicate the strength of the correlation. Significant and marginally significant effects of temperature (T), precipitation (P) and their interaction (TxP) are indicated next to each cluster. Significance codes: *** = p < 0.001; ** = 0.001 < p < 0.01; * = 0.01 < p < 0.05; $ = 0.05 > p > 0.07. For full networks, see Fig S4, for taxonomic and putative functions of each microbial network cluster see Fig S5 and Table S1-S2.

Microbial co-occurrence networks in the rhizosphere were grouped into seven clusters, while in the roots 11 clusters occurred (Fig 2; Fig S4A, S5AB). Temperature, precipitation and their interaction significantly affected microbial clusters in both the rhizosphere and root (Fig 2). All clusters in the rhizosphere network contained both prokaryote and fungal taxa, showing that prokaryotes and fungi responded relatively similar within the rhizosphere compartment (Fig 2; Fig S4A, S5AB). In the root network, various microbial clusters were dominated by either prokaryotes or fungi, indicating that responses of prokaryotes and fungi were in part decoupled within the root compartment (clusters 2, 5, 11; Fig 2; Fig S4B, S5CD). Between the rhizosphere and root network clusters, we observed only three positive correlations, showing that the microbial networks in these two compartments largely showed different patterns and were thus decoupled (Fig 2).

Where possible, we characterised the putative functions that could be performed by each network cluster (Table S1-S2). Most of the rhizosphere network clusters harboured taxa potentially involved in organic matter degradation and N-cycling (Table S1). In contrast, the majority of root clusters harboured taxa potentially involved in plant pathogen attack, the suppression of plant diseases, and the degradation of organic matter (Table S2).

### Temperature, precipitation and their interaction shape plant community composition and soil properties

To understand via which pathways temperature, precipitation, and their interaction shaped the rhizosphere and root microbial communities, we traced their effects through the plant and soil ecosystem using structural equation modelling (SEM). We found that both temperature, precipitation and their interaction had profound effects on plant community composition, which cascaded into affecting local *F. rubra* soil properties (Fig 3).

**Fig. 3.**
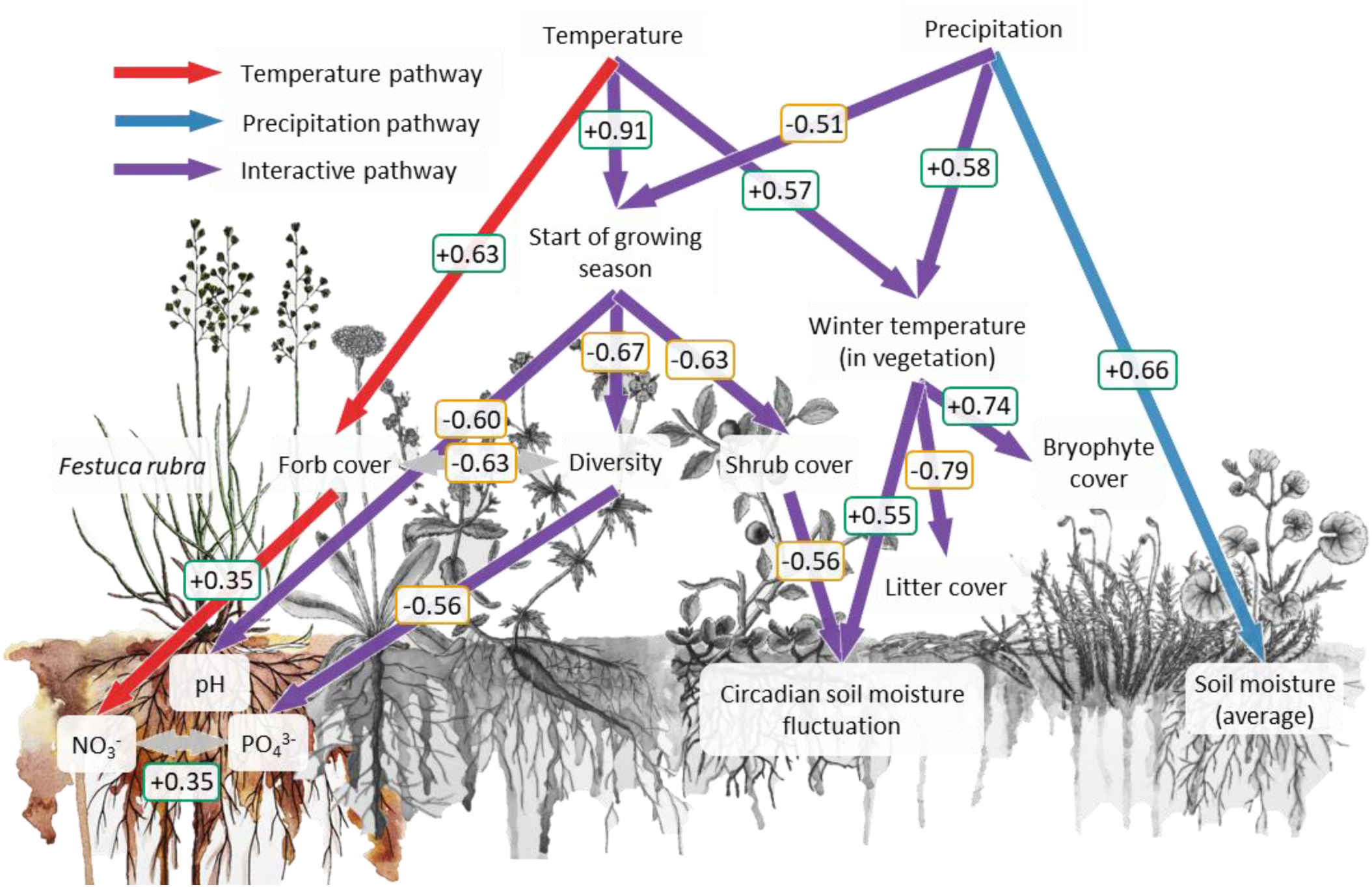
Structural equation model (SEM) showing effects of temperature, precipitation and their interaction on plant community composition, bulk soil properties (circadian soil moisture fluctuation and average soil moisture) and local *Festuca rubra* soil properties (NO_3_^−^, PO_4_^3−^, and pH). Red arrow indicate significant pathways regulated by temperature only, blue arrows indicate pathways regulated by precipitation only, and purple arrows indicate pathways regulated by both temperature and precipitation (interactive pathways) (p < 0.05). Grey arrows indicate significant correlations (p < 0.05). Numbers indicate pathway effect sizes ranging from -1 (negative effects; yellow) to +1 (positive effects; green) (*n* = 12-94 over 12 locations). Absence of arrows indicates that pathways were not significant (p > 0.05).

For both temperature and precipitation, an unique pathway occurred where the two factors did not interact. Precipitation increased the average soil moisture (Fig 3), while temperature increased forb cover and thereby mediated a high availability of NO_3_^−^ in soil below *F. rubra* plants at warm locations (Fig 3).

In contrast, the start of the growing season and average winter temperature in the vegetation canopy were mediated by interacting effects of temperature and precipitation. Specifically, high temperature advanced the start of the growing season, while high precipitation delayed the start of the growing season (Fig 3). The temperature in the vegetation canopy during the winter months was increased by both high temperature and high precipitation (Fig 3).

An early start of the growing season decreased soil pH below *F. rubra* plants, community shrub cover, and plant community diversity. The latter subsequently decreased PO_4_^3-^ availability below *F. rubra* plants, while decreased shrub cover decreased the circadian soil moisture fluctuation at the location (Fig 3). This circadian fluctuation indicates that at low shrub cover, soil moisture dropped during the day time, while at high shrub cover, soil moisture was constant during the day-night cycle (Fig S7). Circadian soil moisture fluctuation was further increased by a high temperature during the winter months. A high winter temperature in the vegetation canopy furthermore increased bryophyte cover, while decreasing litter cover (Fig 3). *F. rubra* cover was not significantly affected by temperature or precipitation (data not shown).

### Interactive pathways of temperature and precipitation were key in shaping microbial co-occurrence networks

We determined via which pathways temperature and precipitation shaped the microbial rhizosphere and root network clusters by combining our SEM model with the microbial network clusters. In the rhizosphere, temperature, precipitation, and their interacting pathways explained between 50-60% of the variation in prokaryote and fungal network clusters (Fig 4AB). Most of this variation was explained by pathways interactively affected by temperature and precipitation (Fig 4AB). The start of the growing season, bryophyte cover and soil pH contributed most strongly to shaping rhizosphere prokaryote and fungal network clusters (Fig 4CD). Especially for fungal network clusters, average soil moisture and forb cover contributed to a lesser, yet significant amount of variation (Fig 4CD).

**Fig. 4.**
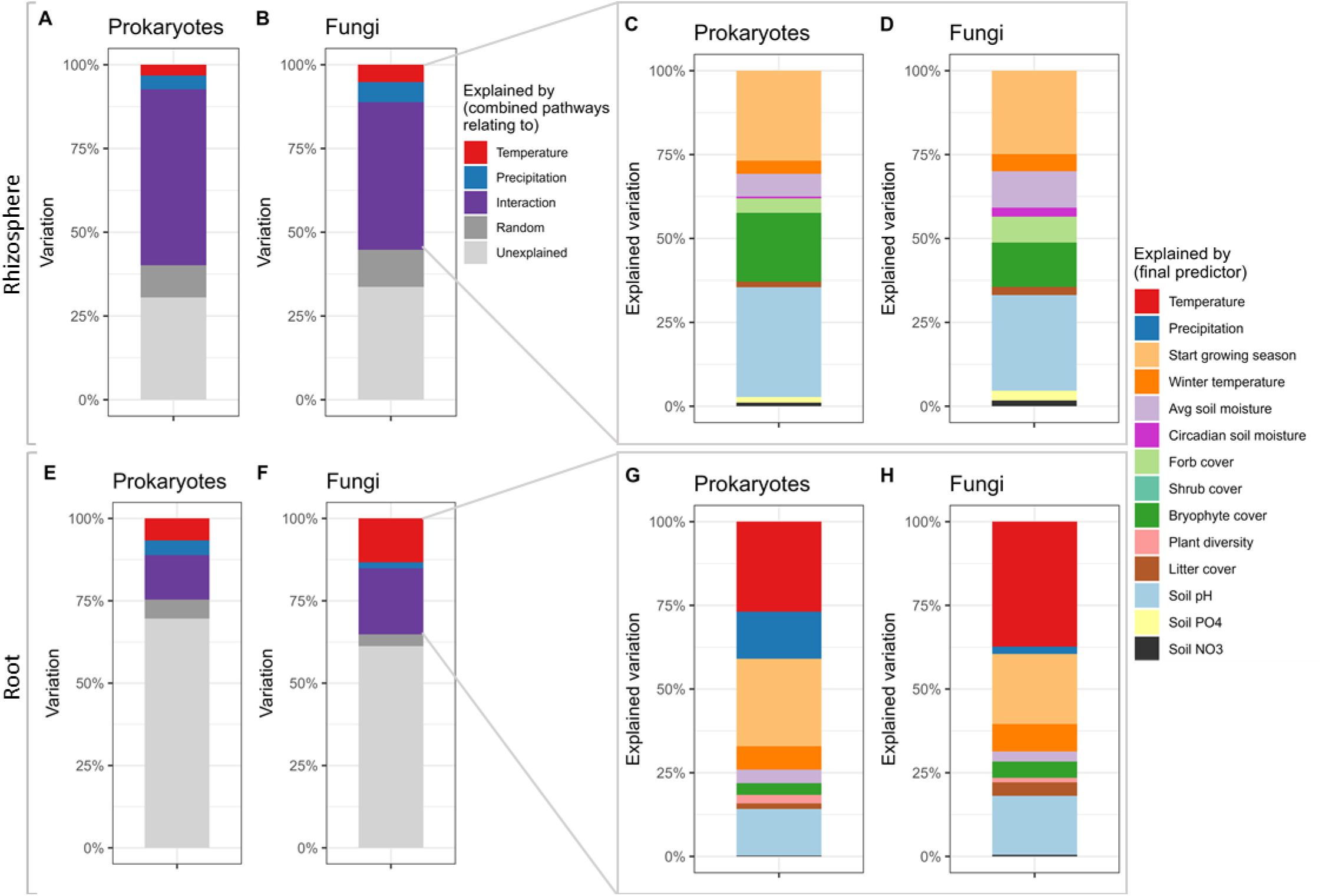
Relative contribution of temperature, precipitation and interacting pathways on (A-D) rhizosphere and (E-H) root-associated microbial communities of *Festuca rubra*. In A, B, E, and F the combined relative contribution of all temperature, precipitation, and interactive pathways in explaining variation in the rhizosphere and root-associated microbial communities are shown. Panels C, D, G, and H zoom in on the explained variation from the respective full overview from A, B, E, or F, separated into the various underlying pathways (see Fig 3 for pathways). Relative contribution of pathways is based on the effect sizes of the structural equation model pathways weighted by the size of the involved microbial rhizosphere or root clusters. Random refers to the relative contribution of within-location variation. N = 94 individuals distributed over 12 locations.

In contrast to the rhizosphere, only 25-35% of variation in root prokaryote and fungal network clusters was explained by temperature, precipitation, and their interaction (Fig 4EF). Especially for fungal root-associated network clusters, pathways shaped by temperature alone played a significant role (Fig 4EF), while for root-associated prokaryote network clusters, direct effects of precipitation explained more variation than in the rhizosphere (Fig 4GH). Yet, similar to the rhizosphere, a large fraction of the explained variation was linked to pathways interactively affected by temperature and precipitation (Fig 4EF). In line with the rhizosphere networks, the start of the growing season and soil pH were important pathways shaping microbial network clusters in the root (Fig 4GH).

### Temperature and precipitation affected putative organic matter degraders, plant pathogens, and plant beneficial microbiota

We identified the microbial clusters affected by the most prominent temperature and precipitation effect pathways (Table S1-S2). In the rhizosphere, an advancement of the start of the growing season and subsequent drop in soil pH was associated with a decrease in relative habitat specialist prokaryotes and fungi. These prokaryotes and fungi were potentially involved in organic matter degradation, including the degradation of recalcitrant C sources (cluster 2 in Table S1). Simultaneously, distinct clusters of microbiota also potentially involved in degrading recalcitrant C sources, were decreased (clusters 3 and 7 in Table S1). The latter group involved microbiota suggested to be facultatively anaerobic (cluster 3 in Table S1). In the root, an early start of the growing season increased mostly putative plant pathogens and bacteria potentially promoting plant growth (clusters 2 and 3 in Table S2). Additionally, via the drop in soil pH with an early start of the growing season, mainly the putative fungal organic matter degrading community shifted in composition (clusters 6, 8, and 10 in Table S2).

In the rhizosphere, bryophyte cover decreased mainly relative habitat generalist microbiota potentially degrading recalcitrant C sources (clusters 2 and 3 in Table S1). In contrast, forb cover, soil moisture, and litter cover increased fungi and relative habitat specialists prokaryotes potentially involved in organic matter degradation (clusters 4, 5, and 7 in Table S1). In the root, direct temperature effects decreased putative organic matter degrading microbiota (cluster 8 in Table S2), while precipitation decreased relative habitat generalist microbiota characterised as putative plant pathogens and beneficial microbiota (cluster 1 in Table S2).

### Contrasting effects of temperature and precipitation on habitat specialisation of microbiota in the rhizosphere and root

Finally, on an overall microbial community scale, we found that increasing temperature decreased and increasing precipitation increased relative habitat specialisation of the prokaryote community. These effects occurred via advances and delays in the start of the growing season and subsequent effects on soil pH (Fig 3, 5AB). Relative habitat specialisation of the rhizosphere fungal community, on the other hand, was decreased with temperature via an increase in forb cover (Fig 3, 5C), and further decreased by both temperature and precipitation via a reduction in litter cover with higher winter temperatures in the vegetation (Fig 3, 5D).

In the root-associated compartment, precipitation directly increased relative habitat specialisation of the prokaryote community (Fig 5E). Relative habitat specialisation of the root-associated fungal community was, similar to the rhizosphere fungal community, decreased by increasing temperature via an increase in forb cover (Fig 3, 5E).

**Fig. 5.**
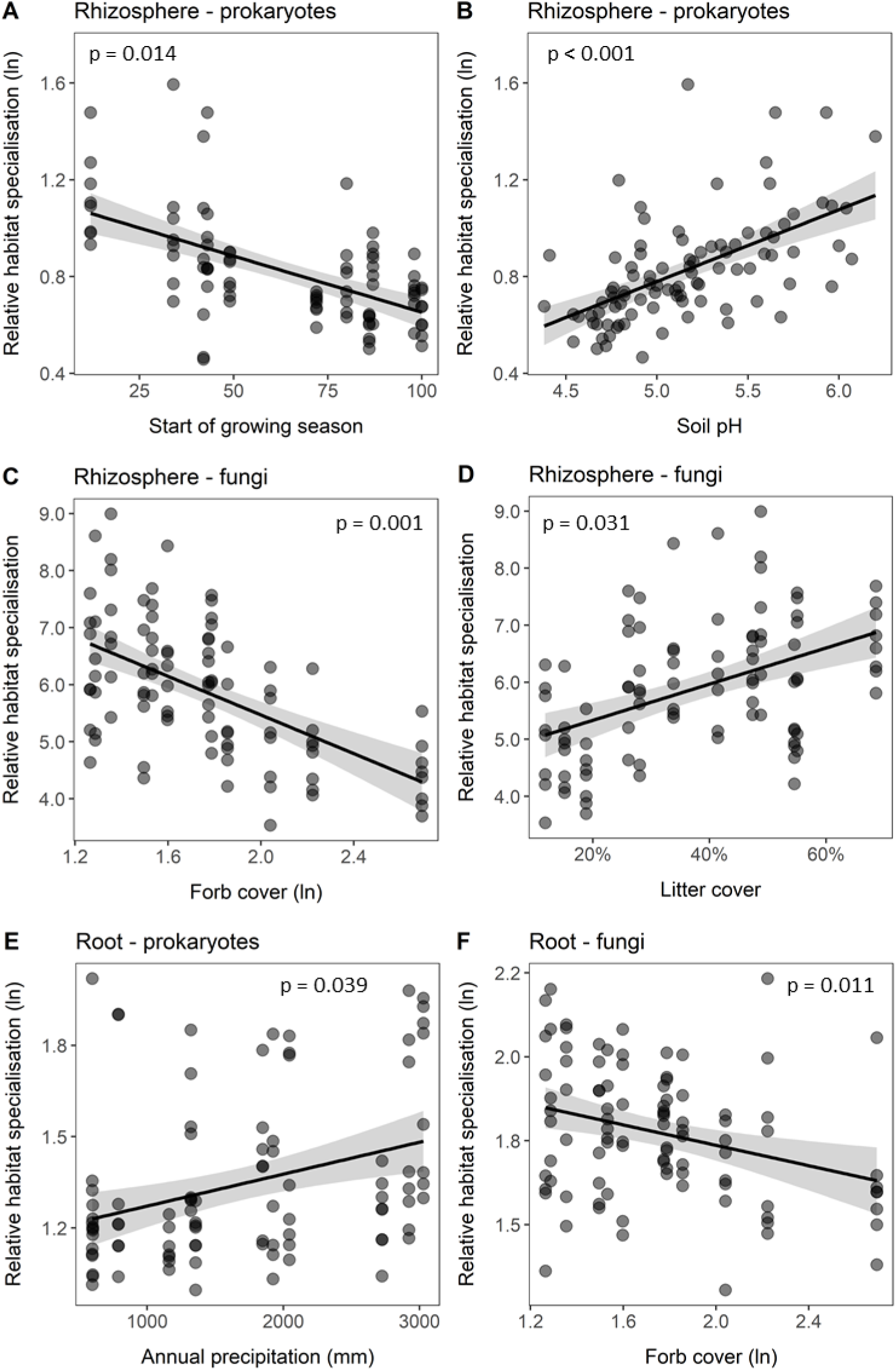
Significant relations between relative habitat specialisation of (A-C) rhizosphere microbiota and (D-F) root-associated microbiota to climate, community vegetation cover and soil properties. The start of the growing season is indicated as the number of days after soil defrosting. Solid lines indicate the mean relation with in grey the 95% confidence interval. Relative habitat specialisation is the community weighted mean of the specialisation index of all ASVs present in a sample. Higher values indicate a greater relative habitat specialisation of the microbial community. Statistical results of the pathways from structural equation models are shown (n = 94 *F. rubra* individuals distributed over 12 locations).

## Discussion

### The start of the growing season mediates interactive effects of temperature and precipitation on plant-soil microbiota interactions

In alpine grassland communities, we tested via which pathways temperature and precipitation shape plant-soil microbiota interactions. Our findings show that the interaction between temperature and precipitation is more critical in shaping the rhizosphere and, to a lesser extent, the root-associated microbiome of *F. rubra* than temperature and precipitation alone. This aligns with Münzbergová et al.^38^, who demonstrated that the interaction of temperature and precipitation is most important in determining the growth strategy of *F. rubra*, such as determining its investment in rhizome and root biomass. Similarly, interactions between temperature and precipitation were found to drive plant community productivity as well as plant species colonisation and extinctions rates within the same climate grid^34,35,39^.

We further found that the interactive effects of temperature and precipitation on plant-microbiota interactions are primarily mediated by changes in the start of the growing season and subsequent effects on soil pH. Specifically, higher temperatures advanced the start of the growing season, leading to a decrease in soil pH. In contrast, higher precipitation delayed the start of the growing season, resulting in an increase in soil pH. These opposing effects likely result from contrasting effects of temperature and precipitation on snow conditions during winter^40^. Higher temperatures lead to quicker snowmelt, advancing the start of the growing season, while increased precipitation leads to greater snow cover, extending snow cover duration, and delaying the start of the growing season^40–42^. Indeed, soil microbial communities undergo marked shifts between winter and summer^31,43^, and snow removal experiments show that an advancement of the start of the growing season significantly reshapes soil microbial communities and reduces soil pH^31–33^. We conclude that in cold environments, the start of the growing season is a critical factor mediating the interactive effects of temperature and precipitation on plant-soil microbiota associations.

Our findings suggest that in locations where both temperature and precipitation are increasing due to climate change, their opposing effects on the start of the growing season moderate changes in plant-soil microbiota associations. In contrast, in locations where temperature is rising and precipitation decreasing, both factors advance the start of the growing season, amplifying each other’s effect on plant-soil microbiota associations. Given the critical role of winter snow cover in these interactions, it is important to note that climate change is causing more winter precipitation to fall as rain^44^. We therefore suggest that in overall cold and wet environments, shifts in plant-soil microbiota associations are particularly accelerated in areas experiencing both warming and drying. Moreover, the transition from snow to rain during winter represent a critical tipping point, likely to further accelerate changes in plant-microbiota interactions.

### An early start of the growing season increased habitat generalist rhizosphere prokaryotes and putative plant pathogens in the root

Changes in the start of the growing season affected rhizosphere and root-associated prokaryotes and fungal communities. An early start of the growing season and subsequent drop in soil pH increased the abundance of habitat generalists prokaryotes in the rhizosphere. Prokaryotes become active after snowmelt when a surge in available N in the soil occurs, and exhibit a fast turnover throughout the season^36,45,46^. Hence, rhizosphere prokaryotes specialised to winter conditions were likely outcompeted as time passed since the start of the growing season, explaining the observed increase in habitat generalists in the rhizosphere.

Many rhizosphere microbiota and root-associated fungi affected by the start of the growing season were likely involved in degrading recalcitrant C sources. From a seasonal perspective, this involvement makes sense, as labile C compounds that accumulate in the soil during autumn and winter are rapidly degraded early in the season following snowmelt. This processes leaves primarily recalcitrant compounds available in summer when we sampled^31,43^. These patterns also suggest high redundancy within the microbial community affected by the start of the growing season, because microbiota performing similar functions appeared to replace one another in response to shifts in the start of the growing season. Consequentially, the functional impact of climate change through shifts in the start of the growing season on rhizosphere microbiota and root-associated fungi may be relatively limited. In contrast, functional redundancy was not evident for the putative plant pathogens and beneficial bacteria in the root that were affected by the start of the growing season. Both groups increased with an early start of the growing season, possibly resulting from a prolonged accumulation time since its onset. Although these findings show that higher temperatures lead to increases in both putative plant pathogens and beneficial microbiota, one of the most commonly observed effects of rising temperatures is an increase in plant disease^47^. Consequentially, positive effects of beneficial microbiota may not offset negative effects of the increase in putative pathogens. We conclude that shifts in the growing season start may play a critical role in increasing plant disease pressure with the ongoing change in climate.

### Temperature and precipitation increased habitat generalist plant-fungal interactions

Where habitat specialisation for the rhizosphere prokaryote community was related to changes in the start of the growing season, different pathways were at play for the rhizosphere and root-associated fungal communities. In both the rhizosphere and root compartments, higher temperatures and increased precipitation led *F. rubra* to associate with more fungal habitat generalists. In both compartments, these patterns resulted from an increase in forb cover with temperature. Additionally, in the rhizosphere, temperature and precipitation increased winter temperatures in the vegetation and subsequently reduced litter cover, which also contributed to an increase in fungal habitat generalists. This increase in generalists suggests a loss of heterogeneity in plant-associated fungal communities as well as an shift towards a greater resistance of the fungal associated communities to environmental change. This is because habitat generalists, with their broad environmental tolerance, are more resilient to environmental change, while habitat specialists, with narrower environmental ranges, are more vulnerable to these changes^48,49^. These shifts are concerning because specialisation in plant-soil microbiota interactions and soil heterogeneity in general are key drivers of plant species coexistence, community diversity and long-term stability^2,8,9,36^. Furthermore, given the high resistance to change of habitat generalists, these observed shifts are likely to be irreversible. We conclude that in cold regions where both temperature and precipitation are increasing, specialist interactions between plants and soil fungi are being lost, which is likely to significantly alter the mechanisms structuring ecosystem dynamics.

Many fungal rhizosphere and root taxa were potentially involved in organic matter degradation, including those related to changes in litter and forb cover. Many fungi specialise in degrading recalcitrant plant litter^43^, which explains why fungal habitat specialists decreased with a reduction in litter cover. Additionally, the decline of specialist fungi with increased plant community forb cover due to rising temperatures is likely related to the capacity of acquisitive forb species to promote the turnover of soil organic matter by exuding labile C compounds in their rhizosphere^50^. This exudation of labile compounds likely favoured fast-growing, habitat generalist microbiota over fungi specialised in degrading recalcitrant organic matter. We conclude that in cold areas experiencing increases in both temperature and precipitation, the degradation of recalcitrant organic matter by plant-associated fungi will increasingly be performed by habitat generalists.

In contrast to the fungal community, the root-associated prokaryote community was increased in habitat specialists with high precipitation. This effect likely resulted from more frequent soil anoxic conditions due to high precipitation, along with associated changes in plant physiology and root traits^27–29,51^. These conditions namely favour specialist prokaryotes that can tolerate or thrive in this specific environment^48^. This selective pressure was likely experienced to a lesser extent by the fungal community, as fungi are generally more resistant to environmental disturbances than prokaryotes^36,45,46^. The prokaryotes affected by precipitation included putative plant pathogens and beneficial bacteria. Our findings therefore suggest that an increase in precipitation in cold environments will result in plant disease dynamics being increasingly regulated by habitat specialist prokaryotes.

### Bryophytes are a key mediator of changes in temperature and precipitation

Bryophytes have been shown to key in regulating soil microclimates and plant species invasion success^34,52^. In line, in the current study, we found that bryophyte cover emerged as a critical factor in decreasing the abundance of microbiota involved in organic matter degradation in the rhizosphere of *F. rubra*. This effect was enhanced by both temperature and precipitation, and was mediated by an increase in winter temperature in the vegetation layer. Climate change increases both summer and winter temperature, while precipitation increases winter temperature in the vegetation by enhancing snow cover, which insulates the vegetation layer^53^. Higher winter temperatures and sustained soil moisture due to snow cover likely increased bryophyte cover^54,55^. Bryophytes then acted as thermal insulators for the soil, reducing diurnal soil temperature fluctuations^52,56^. This higher and more constant soil temperature may have accelerated soil microbial processes, thus decreasing slow-growing rhizosphere microbiota degrading recalcitrant C sources. We conclude that bryophytes play a crucial role in mediating the effects of temperature and precipitation change on plant-soil microbiota interactions.

## Conclusion

We demonstrate that in cold environments, the onset of the growing season is mediated by interactions between temperature and precipitation, and therewith key in shaping interactions between plants and soil microbiota. Temperature and precipitation had opposing effects on the onset of the growing season, likely due to their opposing effects on snow cover duration. These findings highlight the complexity of plant-soil microbial interactions in response to climate and underscore the necessity for local and seasonal monitoring of climatic variables including snow cover to accurately predict ecosystem responses to climate change.

We found that soil pH and bryophyte cover explained a high proportion of the variation in plant rhizosphere and root microbiomes. An early start of the growing season reduced soil pH, while increasing temperature and precipitation both enhanced bryophyte cover. These two variables thus indicate shifts in plant-soil microbiota interactions and may serve as easily measurable indicators for monitoring ecosystem responses to the changing climate.

Our findings further suggest that with rising temperature and precipitation in cold grassland regions, plant-fungal interactions become more tolerant to a wider range of environmental conditions and thus lose specificity, while root prokaryotes gain more specialists. These changes mainly shifted microbial communities performing soil organic matter turnover and putative plant pathogens. These findings indicate that the plant-soil microbiota interactions that structure plant community dynamics are significantly changing with the ongoing climate change^2–9^. We conclude that limiting temperature increase and precipitation change is critical to safeguarding the unique plant-microbiota interactions that shape the functioning of cold climate systems.

## Methods

### Experimental design

The sampling took place in the Vestland climate grid^35,37^. The grid comprises 12 locations in the fjords of southern Norway with factorial combinations of three temperature levels (mean growing season temperatures of 6.5, 8.5 and 10.5 °C) and four precipitation levels (annual precipitation of 600, 1200, 2000 and 2700 mm) (Fig S10). These initial temperature and precipitation levels were calculated based on interpolation of meteorological data from 1961-1990^35,37^. Since 2009, each location in the grid has air and soil temperature sensors and soil moisture sensors installed providing detailed climate information. All 12 locations are natural grassland ecosystems on a calcareous bedrock. The perennial, clonal grass *Festuca rubra* is the only plant species consistently growing at each location and hence was selected as the focal plant species in this study. In the same climate grid, *F. rubra* was found to show both genetic differentiation and plastic responses in its growth traits in relation to temperature, precipitation and their interaction^38,57^. These responses on the plant species level provide a strong basis to suggest consistent changes in the interactions of *F. rubra* with prokaryote and fungal communities within the climate grid as well.

### Sampling

In July 2020, rhizosphere soil and roots of eight individuals of *F. rubra* at each location were collected. The eight individuals per location were selected from a line transect of individuals growing at least 1 m away from each other to avoid sampling of the same *F. rubra* clone. Rhizosphere soil and roots were sampled by taking a soil core of 5 cm in diameter and 5 cm depth at the spot of the *F. rubra* individual. In the field, the plant root system was carefully separated from the bulk soil and only roots attached to the *F. rubra* shoot were collected. Rhizosphere soil was carefully separated from plant roots by brushing the thin layer of soil off the roots and stored at 4 °C. Roots were washed in milliQ water to remove any remaining attached soil participles and loosely attached microbiota. Roots were lightly dried using paper towel and stored on silica gel. Bulk soil for soil chemical analysis was collected from the soil core and stored at 4 °C. In the laboratory, both rhizosphere soil and root tissue were frozen at -80 °C until further analysis.

### Rhizosphere and root microbiome amplicon sequencing

Silicagel dried rhizosphere samples (250 mg each, in duplicates for each sample) and roots (20 mg each, in duplicates for each sample) were homogenized and lysed in PowerBead Tubes (Qiagen, Hilden, Germany) on a Vortex adapter. DNA was extracted using the DNeasy PowerSoil Kit (Qiagen, Germany) according to the manufacturer’s instructions. The fungal internal transcribed spacer of the rDNA (ITS2) was amplified using primers gITS7ngs^58^ and ITS4^59^. The bacterial 16S rRNA gene (V4 region) was amplified from the same DNA extracts using primers 515f and 806r^60^.

PCR amplification was designed in two subsequent reactions: the first PCR (PCR1) was performed with non-barcoded primers, and the second (PCR2) with non-barcoded primers tagged with sample-specific barcodes. The product of the first PCR was used as a DNA template for the second PCR reaction. DNA amplification of fungal sequences was performed in 15 µl (PCR1) and 30 µl (PCR2) reactions as follows: PCR1: 1 x PCR Blue Buffer wo MgCl_2_ (Top-Bio, Vestec, Czech Republic), 2 mM MgCl_2_, 10 µg BSA, 0.2 mM each dNTP, 0.4 µM of each primer, 0.35 U *Taq* DNA Polymerase (Top-Bio, Vestec, Czech Republic) and 20 ng of DNA dissolved in deionized water. PCR2: 1x PCR Blue, 2 mM MgCl_2_, 20 ug BSA, 0.2 mM each dNTP and 0.2 µM of each primer, 0.7 U *Taq* DNA polymerase, 2 µl of PCR1 product. The following thermocycler conditions were used: an initial denaturation step at 94 °C for 5 min, followed by 35 cycles of denaturation (94 °C for 30s), annealing (45 °C for 30s for rhizosphere samples and 50 °C for 30s for root samples) extension (72 °C for 45s), and the final extension at (72 °C for 20min). DNA amplification of bacterial sequences was performed in 15 µl (PCR1) and 30 µl (PCR2) reactions as follows: PCR1: 1x PCR Buffer, 2 mM MgCl_2_, 10 µg BSA, 0.2 mM each dNTP, 0.2 µM of each non-barcoded primer, 0.35 U *Taq* DNA Polymerase and 10 ng of DNA dissolved in deionized water; PCR2: 1x PCR Buffer, 2 mM MgCl_2_, 20 µg BSA, 0.2 mM each dNTP, 0.2 µM of each primer, 0.7 U *Taq* DNA Polymerase and 1 µl of the PCR1 product.

Each DNA sample was extracted in technical duplicates. PCR1 was performed without technical multiplication and PCR2 was performed in technical duplicates to capture maximum diversity from each sample. The pooled tetraplicates were purified by using QIAquick PCR Purification Kit (Qiagen, Hilden, Germany) according to the manufacturer’s protocol. DNA quantification was assessed by the Qubit 2.0 Fluorometer (Thermo Fisher Scientific). The purified amplicons were pooled in equimolar ratios. Two rhizosphere and two root negative PCR controls (ddH_2_O instead of a template) were included in the described workflow. The final amplicon library was sequenced on an Illumina MiSeq instrument (2 × 250 bp) (SEQme, Dobříš, Czech Republic).

### 16S and ITS bioinformatics

In total, we sequenced 96 rhizosphere and 96 root samples with two negative controls each. Raw reads were processed using the pipeline SEED2 ver. 2.1.1b^61^ in which reads were demultiplexed (no mismatch allowed in the tag sequences). Low-quality sequences with a mean quality score < 30 were discarded, including reads with non-matching tags. Primers and barcode sequences were cut from the reads.

All following bioinformatics on demultiplexed raw FASTQ files were analysed using the DADA2 pipeline for prokaryote 16S and fungal ITS sequences in R version 4.1.2^62,63^. Both 16S and ITS sequences were quality trimmed and filtered according to the standard DADA2 protocols for 2 x 250 bp reads. Increasing the maxEE quality filter did not improve the quality filtering of ITS sequences^64^. After filtering and trimming, sequences were inferred based on a parametric error model, after which chimeric sequences and ASVs occurring in the blank samples were removed as these were likely contaminants. Taxonomic information was obtained using the SILVA (v138) and UNITE (v8.2) databases for 16S and ITS, respectively^65,66^. Non-prokaryote and non-fungal hits were removed as well as mitochondrial and chloroplast hits. The procedure resulted in 35067 unique 16S ASVs and 8881 unique ITS ASVs. These were represented by 4098759 and 4611230 total reads for the 16S and ITS amplicons, respectively. ASVs with in total < 100 reads were removed as well as one sample with in total < 1000 reads, and an outlier likely originating from a plant that was likely not *Festuca rubra*.

### Microbial community calculations

Read counts were normalised using centered log ratio (clr) transformation and visualised using PCA with the phyloseq package^67^. Significant separation between locations was tested with PERMANOVA using *adonis* of the vegan package^68^. A passive overlay of temperature and precipitation gradients were created using *envfit* of the vegan package^68^. Shannon-diversity was calculated based on multiple rarefaction (1000 iterations) using the metagMisc package^69^.

For each ASV, we calculated a specialisation index (SI) for the rhizosphere and root associated compartments each following Chen et al^48^. SI of the *i*th ASV was calculated as the coefficient of variation minus a correction for under-sampling of rare ASVs:

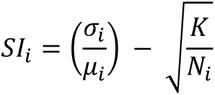

with *σ*_*i*_ being the standard deviation of the reads of the *i*th ASV across all samples, *μ*_*i*_ being the mean of the reads of the *i*th ASV across all samples, K the number of habitat classes (12 locations) and N the total number of reads of the *i*th ASV across all samples. Calculations were based on rarefied read abundances (rarefied to the smallest sample size: 3766 and 1828 for 16S and ITS samples, respectively). Average SI of each sample was calculated as the community weighted mean of the SI of all ASVs present in the sample.

### Microbial co-occurrence networks and cluster characteristics

We constructed microbial co-occurrence networks for the rhizosphere and root compartment each using the SPIEC-EASI method^70^. Prokaryotes and fungi were incorporated together in each of the two networks^71^. We first excluded rare ASVs < 100 reads in total and ASVs that occurred in < 6 samples. Co-occurrence networks were then calculated based on clr-transformed read counts and the neighbourhood selection method with optimised stability parameters based on the StARS selection procedure (*n* = 94 *Festuca* plants; threshold 0.05, nlambda 45 with 300 replications)^72^. Similarly responding ASVs in each network were clustered using the Spin-glass algorithm of the igraph package, which includes both positive and negative edges with similar importance and unlimited clusters^73–76^. For each cluster in each network, we summed the relative read counts of the ASVs per sample and tested for significant correlations between clusters using linear mixed effects models using lme of the nlme package^77^. Location was incorporated as a random effect in these models. Relative read counts of clusters were ln- or sqrt-transformed in case model residuals did not follow a normal distribution. P-values were adjusted for multiple comparisons using the Bonferroni correction^78^.

For each microbial cluster in the rhizosphere and root network we calculated a set of characteristics to infer its putative function. We calculated the average relative abundance of microbial phyla, classes, orders and families and, where possible, inferred putative characteristics of the cluster based on knowledge obtained from literature on the present microbial taxa (Table S1-S2). For each cluster, we calculated the average relative size and the average relative abundance of prokaryotes and fungi based on relative read counts (Table S3-S4).

We also calculated the number of unique prokaryote and fungal ASVs present in each cluster and defined whether the cluster held relative habitat generalists or specialists (Table S3-S4). For this, we calculated the community weighted mean of the SI of each sample in each cluster. Relative habitat generalist clusters were defined as clusters below the community-wide mean SI, while relative habitat specialists were defined as clusters above the community-wide mean SI^48^. Clusters with an overlap in interquartile range of SIs with the community-wide mean SI were defined as unspecified (Fig S8).

### Soil property measurements

Soil collected from below *F. rubra* was sieved on a 2 mm mesh and thoroughly mixed. Plant-available nitrogen (N) (mg kg^−1^ dry soil) was measured by extracting N compounds using 50 mL of 0.5 M K_2_SO_4_ with 5 g of fresh soil. The mixture was shaken for 2 h after which the soil was filtered out. NO_3_^−^, NH_4_^+^ and NO ^−^ concentrations were determined in the filtrate by Flow Injection Analysis (QuickChem 8000 FIA; Lachat Instruments, Loveland, CO, USA). To determine plant-available phosphorus (P), 5 g air dried soil was extracted with 50 mL of 1 M NaHCO_3_. The solution was adjusted to pH 8.5 by adding activated carbon to eliminate discoloration resulting from humic acid release. The solution was shaken for 30 min, soil was filtered out, and available P was determined in the filtrate by the Olsen photometric method (ATI Unicam UV 400/VIS Spectrophotometer at 630 nm)^79^. Soil pH was measured by shaking 5 g air dried soil with 25 mL deionised water for 30 min and measuring pH in the filtrate (WTW Multilab 540; Xylem Analytics, Weilheim, Germany). Soil total N and C content were determined using combustion analyses (FLASH 2000 CHNS/O Analyzer; Thermo Fisher Scientific, Waltham, MA, USA) on dried soil ground to <0.1 mm particle size.

### Start of the growing season, temperature and soil moisture calculations

At each location, the start of the growing season was calculated as the number of days between the first date in 2020 when soil temperatures rose above 2.5 °C and our sampling date in summer. Mean winter temperature in the vegetation layer was determined as the 12-year mean temperature at 30 cm above the soil during the winter period from January until March (soils began to thaw in April for the first time since winter). For each soil moisture sensor at each location, we calculated the 12-year average daily mean soil moisture from 2009 until 2020. These values were averaged values across the two sensors per location. Circadian soil moisture fluctuation was determined by averaging soil moisture levels for each hour across all years and calculating the coefficient of variation over this 24 hour period (Fig S4). In cases where soil moisture or temperature sensors failed for more than 5 months in a given year, that year was excluded from the calculation.

We tested whether the 12-year mean values for average winter temperature in the vegetation, average soil moisture, and circadian soil moisture fluctuation showed significantly different patterns compared to the values calculated for the year 2020 alone. Since no significant differences were found, we chose to use the 12-year mean values, as these provided averages with lower error rates (data not shown).

### Plant community composition

Plant species cover data was obtained from control plots of a climate change transplant experiment by Vandvik et al^34^ and Lynn et al^80^. At each location, five 25 x 25 cm plots provided plant community cover data on a plant species level, of which we used data from 2019 only. For each plant species occurring in the plot, the cover was visually estimated and plant community diversity was calculated for each location using the Shannon diversity index and vegan package^68^. Cover estimates were summed per functional plant group (forbs, grasses excluding *F. rubra*, legumes, shrubs, pteridophytes, hemiparasites, and sedges and rushes together) and *Festuca rubra*. Total bryophyte cover and soil litter cover were obtained from measurements in the same plots obtained in 2013, when a detailed study of bryophytes and litter cover in the plots was conducted (Pascal et al. unpublished)^34,35^. We consider the combination of data from different years to be justified, because of the generally slow growth of bryophytes in cold climates and the minimal shifts in plant community composition that were previously observed over time^34,80^.

### Structural equation models

We hypothesised that temperature and precipitation affected plant-soil microbiota interactions directly, were mediated by changes in the start of the growing season, via plant community composition and/or through soil abiotic properties. We used structural equation models (SEM) to separate these different pathways that may play a role in mediating temperature and precipitation effects (Fig 2). All SEMs were fit using piecewiseSEM^81^ and lme of the nlme package^77^ with location as random effect. Model fit was assessed using directional separation tests (d-sep) based on Fisher’s C statistics with models being accepted if p > 0.05. We simplified our models using a backward stepwise elimination procedure for which we consecutively removed pathways with the highest p-value^82^. Endogenous variables were allowed to drop from the model in case effects were not significant (p > 0.05). The model with the lowest Akaike information criterion (corrected for small sample sizes; AICc) was selected as the best fit base model.

Following in ‘t Zandt et al.^36^, we first calculated a base model defining the significant pathways via which temperature and precipitation affect the start of the growing season, winter temperature in the vegetation, soil moisture variables, plant community composition, and soil abiotic properties. Next, each microbial network cluster (summed relative reads per rhizosphere and root microbial network cluster from the calculated co-occurrence networks; 16S and ITS ASVs separately), as well as alpha-diversity indices and average SI of the microbial communities were ran through the SEM model as the final parameter to be estimated. Per run, one microbial parameter was considered, which could be affected either directly by temperature and precipitation and/or indirectly via the start of the growing season, plant community composition and/or soil abiotic properties. Each run, a backward stepwise elimination procedure to consecutively remove non-significant pathways was followed in the same way as performed for the base part of the SEM. All microbial variables not following a normal distribution were ln- or sqrt-transformed.

### Calculation of explained variation microbial co-occurrence networks

In total, we created 34 unique SEM models based on the microbial co-occurrence networks: 7 for 16S rhizosphere network clusters, 7 for ITS rhizosphere network clusters, 11 for 16S root network clusters, and 9 for ITS root network clusters. From each mode, we extracted the effect sizes of each significant pathway (p < 0.05). For each significant pathway, we calculated composite effect sizes by multiplying the effect sizes between intermediate variables, where applicable. We summed the effect sizes of all significant pathways affecting a microbial network cluster for each climate, plant, and soil variable^36^. These were then scaled to the relative size of the involved network clusters and the percentage of explained variation of the SEM for the involved network cluster.

## Data availability

All raw data generated in this study will be deposited in the Zenodo digital data repository upon acceptance of the manuscript. The raw microbial sequencing data generated in this study will be deposited in the NCBI SRA database. Fungal trait data is publicly available via the FungalTrait database^83^.

## Code availability

All R scripts will be made publicly available via Github and the Zenodo digital data repository upon acceptance of the manuscript.

## Acknowledgements

We are grateful to Linn Krüger, Silje Östman and Susanne Berthelsen for field site maintenance, Siri Lie Olsen and Pascale Michel for assistance with vegetation sampling, Věroslava Hadincová for help with the *Festuca rubra* soil and microbiome field sampling, Andrea Jarošová for help with preparing samples for amplicon sequencing, Zuzana Kolaříková for help with bioinformatics, Kerry Ryan for helpful comments on the manuscript, and Natalie Oram for statistical and microbial network discussions. D.Z. is grateful to Michael Bahn and Fiona Brennan for their support and hosting her at their groups. Z.M., M.Š., A.F. and D.Z. acknowledge funding from The Czech Science Foundation (GAČR 22-00761S), which funded all the data collection for this paper except for the vegetation and climate data, long-term research development project RVO 67985939 of the Czech Academy of Sciences, and the Ministry of Education of the Czech Republic (MŠMT). V.V. acknowledges funding from The Norwegian Research Council (184912, 244525, 315249). D.Z. was supported by a PPLZ Postdoctoral Fellowship awarded by the Czech Academy of Sciences.

## Author contributions

Z.M. obtained funding for and set up the *Festuca rubra* microbiome sampling, V.V. obtained the funding for, designed, and maintained the field sites and transplant experiments; V.V. and K.K. set up the field sites and transplant experiment and led the vegetation sampling; Z.M. and A.F. led the rhizosphere, root and soil sampling campaign; M.Š. led the rhizosphere and root DNA extractions including library preparations for amplicon sequencing; D.Z. performed microbial bioinformatics with input from M.H.Z.; D.Z. analysed microbial, soil and plant data; D.Z. wrote the manuscript, all others reviewed and commented. All authors contributed critically to the manuscript and gave final approval for publication.

## References

1. Bardgett, R. D. & Caruso, T. Soil microbial community responses to climate extremes: resistance, resilience and transitions to alternative states. Philosophical Transactions of the Royal Society B: Biological Sciences 375, 20190112 (2020).

2. van der Putten, W. H., Bradford, M. A., Brinkman, P. E., van de Voorde, T. F. J. & Veen, G. F. Where, when and how plant-soil feedback matters in a changing world. Funct Ecol 30, 1109–1121 (2016).

3. Bever, J. D., Platt, T. G. & Morton, E. R. Microbial population and community dynamics on plant roots and their feedbacks on plant communities. Annu Rev Microbiol 66, 265–283 (2012).

4. Semchenko, M. et al. Soil biota and chemical interactions promote co-existence in co-evolved grassland communities. Journal of Ecology 107, 2611–2622 (2019).

5. Aldorfová, A., Knobová, P. & Münzbergová, Z. Plant–soil feedback contributes to predicting plant invasiveness of 68 alien plant species differing in invasive status. Oikos 129, 1257–1270 (2020).

6. Kardol, P., Bezemer, T. M. & van der Putten, W. H. Temporal variation in plant-soil feedback controls succession. Ecol Lett 9, 1080–1088 (2006).

7. Engelkes, T. et al. Successful range-expanding plants experience less above-ground and below-ground enemy impact. Nature 456, 946–948 (2008).

8. in ‘t Zandt, D., et al. Plant life-history traits rather than soil legacies determine colonisation of soil patches in a multi-species grassland. Journal of Ecology 110, 889–901 (2022).

9. in ‘t Zandt, D., Herben, T., van den Brink, A., Visser, E. J. W. & de Kroon, H. Species abundance fluctuations over 31 years are associated with plant–soil feedback in a species-rich mountain meadow. Journal of Ecology 109, 1511–1523 (2021).

10. IPCC. Climate Change 2021: The Physical Science Basis. Contribution of Working Group I to the Sixth Assessment Report of the Intergovernmental Panel on Climate Change. (Cambridge University Press, 2021).

11. Philippot, L., Raaijmakers, J. M., Lemanceau, P. & van der Putten, W. H. Going back to the roots: the microbial ecology of the rhizosphere. Nat Rev Microbiol 11, 789–99 (2013).

12. Dini-Andreote, F. Endophytes: the second layer of plant defense. Trends Plant Sci 25, 319–322 (2020).

13. Trivedi, P., Batista, B. D., Bazany, K. E. & Singh, B. K. Plant–microbiome interactions under a changing world: responses, consequences and perspectives. New Phytologist 234, 1951–1959 (2022).

14. Ferrari, A., Hagedorn, F. & Niklaus, P. A. Disentangling effects of air and soil temperature on C allocation in cold environments: a 14C pulse-labelling study with two plant species. Ecol Evol 8, 7778–7789 (2018).

15. Medlyn, B. E. et al. Temperature response of parameters of a biochemically based model of photosynthesis. II. A review of experimental data. Plant Cell Environ 25, 1167–1179 (2002).

16. Wang, N. et al. Effects of climate warming on carbon fluxes in grasslands— a global meta-analysis. Glob Chang Biol 25, 1839–1851 (2019).

17. Bengtson, P., Barker, J. & Grayston, S. J. Evidence of a strong coupling between root exudation, C and N availability, and stimulated SOM decomposition caused by rhizosphere priming effects. Ecol Evol 2, 1843–1852 (2012).

18. Zhang, X. et al. Rhizosphere hotspots: root hairs and warming control microbial efficiency, carbon utilization and energy production. Soil Biol Biochem 148, (2020).

19. Zhu, B. & Cheng, W. Rhizosphere priming effect increases the temperature sensitivity of soil organic matter decomposition. Glob Chang Biol 17, 2172–2183 (2011).

20. Keuper, F. et al. Carbon loss from northern circumpolar permafrost soils amplified by rhizosphere priming. Nat Geosci 13, 560–565 (2020).

21. Leff, J. W. et al. Consistent responses of soil microbial communities to elevated nutrient inputs in grasslands across the globe. Proceedings of the National Academy of Sciences 112, 10967–10972 (2015).

22. Wang, C. & Kuzyakov, Y. Mechanisms and implications of bacterial–fungal competition for soil resources. ISME J 18, (2024).

23. Crowther, T. W. et al. Quantifying global soil carbon losses in response to warming. Nature 540, 104–108 (2016).

24. Kirschbaum, M. U. F. Will changes in soil organic carbon act as a positive or negative feedback on global warming? Biogeochemistry 48, 21–51 (2000).

25. Lin, Y. et al. Differential effects of redox conditions on the decomposition of litter and soil organic matter. Biogeochemistry 154, 1–15 (2021).

26. Tiedje, J. M., Sexstone, A. J., Parkin, T. B. & Revsbech, N. P. Anaerobic processes in soil. Plant Soil 76, 197–212 (1984).

27. Oram, N. J. et al. Plant traits of grass and legume species for flood resilience and N2O mitigation. Funct Ecol 35, 2205–2218 (2021).

28. Parent, C., Capelli, N., Berger, A., Crèvecoeur, M. & Dat, J. F. An overview of plant responses to soil waterlogging. Plant Stress 20–27 (2008).

29. Oram, N. J. et al. Plant community flood resilience in intensively managed grasslands and the role of the plant economic spectrum. Journal of Applied Ecology 57, 1524–1534 (2020).

30. Rixen, C., et al. Winters are changing: snow effects on Arctic and alpine tundra ecosystems. Arct Sci 8, 572–608 (2022).

31. Broadbent, A. A. D. et al. Climate change alters temporal dynamics of alpine soil microbial functioning and biogeochemical cycling via earlier snowmelt. ISME J 15, 2264–2275 (2021).

32. Gavazov, K. et al. Winter ecology of a subalpine grassland: effects of snow removal on soil respiration, microbial structure and function. Science of the Total Environment 590–591, 316–324 (2017).

33. Broadbent, A. A. D. et al. Climate change disrupts the seasonal coupling of plant and soil microbial nutrient cycling in an alpine ecosystem. Glob Chang Biol 30, 1–14 (2024).

34. Vandvik, V. et al. Biotic rescaling reveals importance of species interactions for variation in biodiversity responses to climate change. Proceedings of the National Academy of Sciences 117, 22858–22865 (2020).

35. Klanderud, K., Vandvik, V. & Goldberg, D. The importance of biotic vs. abiotic drivers of local plant community composition along regional bioclimatic gradients. PLoS One 10, e0130205 (2015).

36. in ‘t Zandt, D., Kolaříková, Z., Cajthaml, T. & Münzbergová, Z. Plant community stability is associated with a decoupling of prokaryote and fungal soil networks. Nat Commun 14, 3736 (2023).

37. Klanderud, K., Meineri, E., Töpper, J., Michel, P. & Vandvik, V. Biotic interaction effects on seedling recruitment along bioclimatic gradients: testing the stress-gradient hypothesis. Journal of Vegetation Science 28, 347–356 (2017).

38. Münzbergová, Z., Hadincová, V., Skálová, H. & Vandvik, V. Genetic differentiation and plasticity interact along temperature and precipitation gradients to determine plant performance under climate change. Journal of Ecology 105, 1358–1373 (2017).

39. Meineri, E., Spindelböck, J. & Vandvik, V. Seedling emergence responds to both seed source and recruitment site climates: a climate change experiment combining transplant and gradient approaches. Plant Ecol 214, 607–619 (2013).

40. Yu, Z. et al. Effects of seasonal snow on the growing season of temperate vegetation in China. Glob Chang Biol 19, 2182–2195 (2013).

41. Kumar, M., Wang, R. & Link, T. E. Effects of more extreme precipitation regimes on maximum seasonal snow water equivalent. Geophys Res Lett 39, 1–6 (2012).

42. Rebetez, M. Theoretical and applied climatology seasonal relationship between temperature, precipitation and snow cover in a mountainous region. Theor. Appl. Climatol 54, 99–106 (1996).

43. Bardgett, R., Bowman, W., Kaufmann, R. & Schmidt, S. A temporal approach to linking aboveground and belowground ecology. Trends Ecol Evol 20, 634–641 (2005).

44. Mankin, J. S. & Diffenbaugh, N. S. Influence of temperature and precipitation variability on near-term snow trends. Clim Dyn 45, 1099–1116 (2015).

45. de Vries, F. T. et al. Soil bacterial networks are less stable under drought than fungal networks. Nat Commun 9, 3033 (2018).

46. Oram, N. J. et al. Drought intensity alters productivity, carbon allocation and plant nitrogen uptake in fast versus slow grassland communities. Journal of Ecology 111, 1681–1699 (2023).

47. Singh, B. K. et al. Climate change impacts on plant pathogens, food security and paths forward. Nat Rev Microbiol 21, 640–656 (2023).

48. Chen, Y.-J. et al. Metabolic flexibility allows bacterial habitat generalists to become dominant in a frequently disturbed ecosystem. ISME J 15, 2986–3004 (2021).

49. Arraiano-Castilho, R. et al. Habitat specialisation controls ectomycorrhizal fungi above the treeline in the European Alps. New Phytologist 229, 2901–2916 (2021).

50. Henneron, L., Kardol, P., Wardle, D. A., Cros, C. & Fontaine, S. Rhizosphere control of soil nitrogen cycling: a key component of plant economic strategies. New Phytologist 228, 1269–1282 (2020).

51. Hartman, S. et al. Ethylene-mediated nitric oxide depletion pre-adapts plants to hypoxia stress. Nat Commun 10, 4020 (2019).

52. Jaroszynska, F. et al. Bryophytes dominate plant regulation of soil microclimate in alpine grasslands. Oikos 2023, 1–14 (2023).

53. Liu, Y., Hansen, B. U., Elberling, B. & Westergaard-Nielsen, A. Snow depth and the associated offset in ground temperatures in a landscape manipulated with snow-fences. Geoderma 438, 116632 (2023).

54. Górski, P., Gądek, B. & Gąbka, M. Snow as a parameter of bryophyte niche partitioning in snow-beds of the Tatra Mountains (Western Carpathians). Ecol Indic 113, 106258 (2020).

55. Cooper, E. J., Little, C. J., Pilsbacher, A. K. & Mörsdorf, M. A. Disappearing green: shrubs decline and bryophytes increase with nine years of increased snow accumulation in the High Arctic. Journal of Vegetation Science 30, 857–867 (2019).

56. Lindo, Z., Nilsson, M. & Gundale, M. J. Bryophyte-cyanobacteria associations as regulators of the northern latitude carbon balance in response to global change. Glob Chang Biol 19, 2022–2035 (2013).

57. Kosová, V., Hájek, T., Hadincová, V. & Münzbergová, Z. The importance of ecophysiological traits in response of *Festuca rubra* to changing climate. Physiol Plant 174, e13608 (2022).

58. Ihrmark, K. et al. New primers to amplify the fungal ITS2 region - evaluation by 454-sequencing of artificial and natural communities. FEMS Microbiol Ecol 82, 666–677 (2012).

59. White, T., Bruns, T., Lee, S. & Taylor, J. Amplification and direct sequencing of fungal ribosomal RNA genes for phylogenetics. in PCR protocols, a guide to methods and applications (ed. White, T.) 315–322 (Academic Press, 1990).

60. Caporaso, J. G. et al. Global patterns of 16S rRNA diversity at a depth of millions of sequences per sample. Proceedings of the National Academy of Sciences 108, 4516–4522 (2011).

61. Větrovský, T., Baldrian, P. & Morais, D. SEED 2: A user-friendly platform for amplicon high-throughput sequencing data analyses. Bioinformatics 34, 2292–2294 (2018).

62. Callahan, B. J. et al. DADA2: High-resolution sample inference from Illumina amplicon data. Nat Methods 13, 581–583 (2016).

63. R Core Team. R: a language and environment for statistical computing. (2021).

64. Rolling, T., Zhai, B., Frame, J., Hohl, T. M. & Taur, Y. Customization of a DADA2-based pipeline for fungal internal transcribed spacer 1 (ITS1) amplicon data sets. JCI Insight 7, e151663 (2022).

65. Nilsson, R. H. et al. The UNITE database for molecular identification of fungi: handling dark taxa and parallel taxonomic classifications. Nucleic Acids Res 47, D259–D264 (2019).

66. Quast, C. et al. The SILVA ribosomal RNA gene database project: Improved data processing and web-based tools. Nucleic Acids Res 41, 590–596 (2013).

67. McMurdie, P. J. & Holmes, S. Phyloseq: an R package for reproducible interactive analysis and graphics of microbiome census data. PLoS One 8, e61217 (2013).

68. Oksanen, J., et al. vegan: community ecology package. https://cran.r-project.org/package=vegan (2018).

69. Mikryukov, V. vmikk/metagMisc: Miscellaneous functions for metagenomic analysis (v.0.0.0.9000). (2021) doi:10.5281/zenodo.571403.

70. Kurtz, Z. D. et al. Sparse and compositionally robust inference of microbial ecological networks. PLoS Comput Biol 11, 1–25 (2015).

71. Tipton, L. et al. Fungi stabilize connectivity in the lung and skin microbial ecosystems. Microbiome 6, 1–14 (2018).

72. Liu, H., Roeder, K. & Wasserman, L. Stability Approach to Regularization Selection (StARS) for high dimensional graphical models. Adv Neural Inf Process Syst 24, 1432–1440 (2010).

73. Newman, M. E. J. & Girvan, M. Finding and evaluating community structure in networks. Phys Rev E Stat Nonlin Soft Matter Phys 69, 1–15 (2004).

74. Reichardt, J. & Bornholdt, S. Statistical mechanics of community detection. Phys Rev E Stat Nonlin Soft Matter Phys 74, 1–16 (2006).

75. The igraph Core Team. igraph - The network analysis package. (2020).

76. Traag, V. A. & Bruggeman, J. Community detection in networks with positive and negative links. Phys Rev E Stat Nonlin Soft Matter Phys 80, 1–6 (2009).

77. Pinheiro, J., Bates, D., DebRoy, S., Sarkar, D. & R Core Team. nlme: linear and nonlinear mixed effects models. https://cran.r-project.org/package=nlme (2019).

78. Bonferroni, C. E. Teoria statistica delle classi e calcolo delle probabilità. Pubblicazioni del R Istituto Superiore di Scienze Economiche e Commerciali di Firenze (1936).

79. Olsen, S., Cole, C., Watanabe, F. & Dean, L. Estimation of available phosphorus in soils by extraction with sodium bicarbonate. in *USDA Circular Nr 939* (US Gov. Print. Office, Washington, DC, USA, 1954).

80. Lynn, J. S., Klanderud, K., Telford, R. J., Goldberg, D. E. & Vandvik, V. Macroecological context predicts species’ responses to climate warming. Glob Chang Biol 27, 2088–2101 (2021).

81. Lefcheck, J. S. piecewiseSEM: piecewise structural equation modelling in R for ecology, evolution, and systematics. Methods Ecol Evol 7, 573–579 (2016).

82. in ‘t Zandt, D., et al. Local soil legacy effects in a multispecies grassland community are underlain by root foraging and soil nutrient availability. Journal of Ecology 108, 2243–2255 (2020).

83. Põlme, S. et al. FungalTraits: a user-friendly traits database of fungi and fungus-like stramenopiles. Fungal Divers 105, 1–16 (2020).

