## Supplementary material for "Temperature and precipitation interactively shape the plant microbiome by regulating the start of the growing season"

### Supplementary Figures

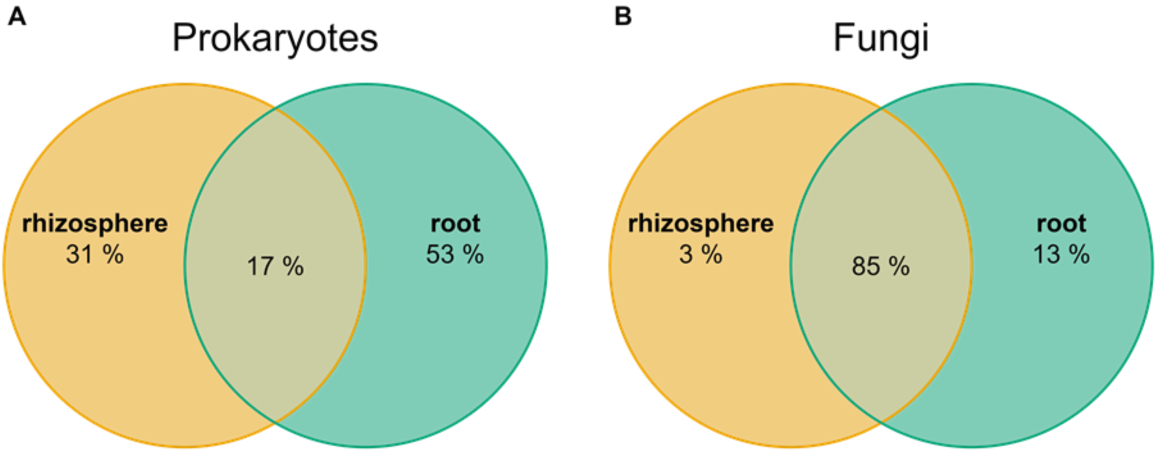


**Fig. S1** Venn diagram showing the overlap in (A) prokaryote and (B) fungal ASVs between the rhizosphere (yellow) and root (green) compartment of *Festuca rubra*. Percentages indicate the number of ASVs in each compartment weighted by the ASVs relative abundance (n = 94 per rhizosphere and root compartment).

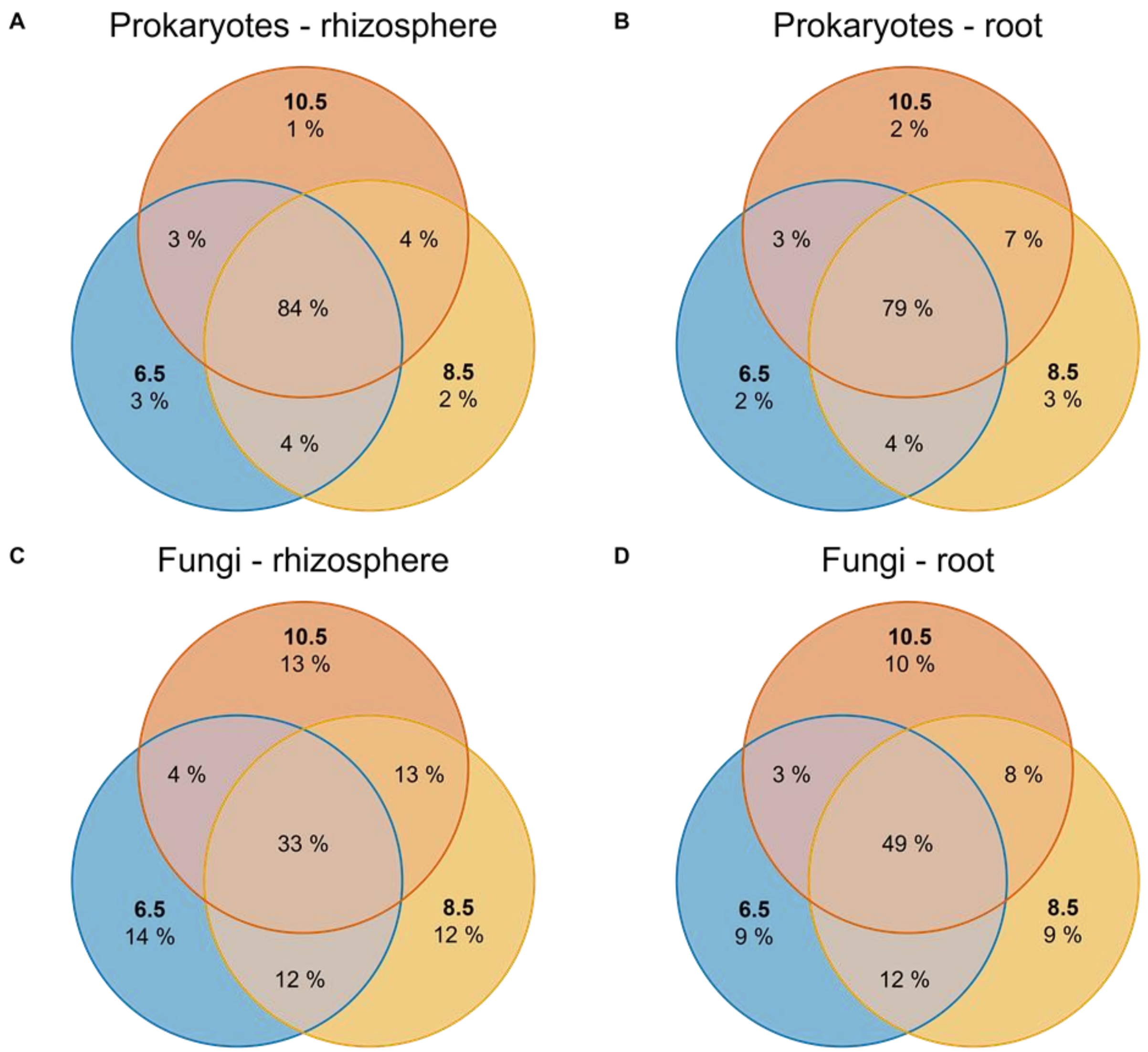


**Fig. S2** Venn diagram showing the overlap in (A) prokaryote ASVs in the rhizosphere, (B) prokaryote ASVs associated to the root, (C) fungal ASVs in the rhizosphere, and (D) fungal ASVs associated to the root between the three levels of average summer temperature at locations: 6.5 (blue), 8.5 (yellow), and 10.5 (orange). Percentages indicate the number of ASVs in each compartment weighted by the ASVs relative abundance (n = 94 per rhizosphere and root compartment).

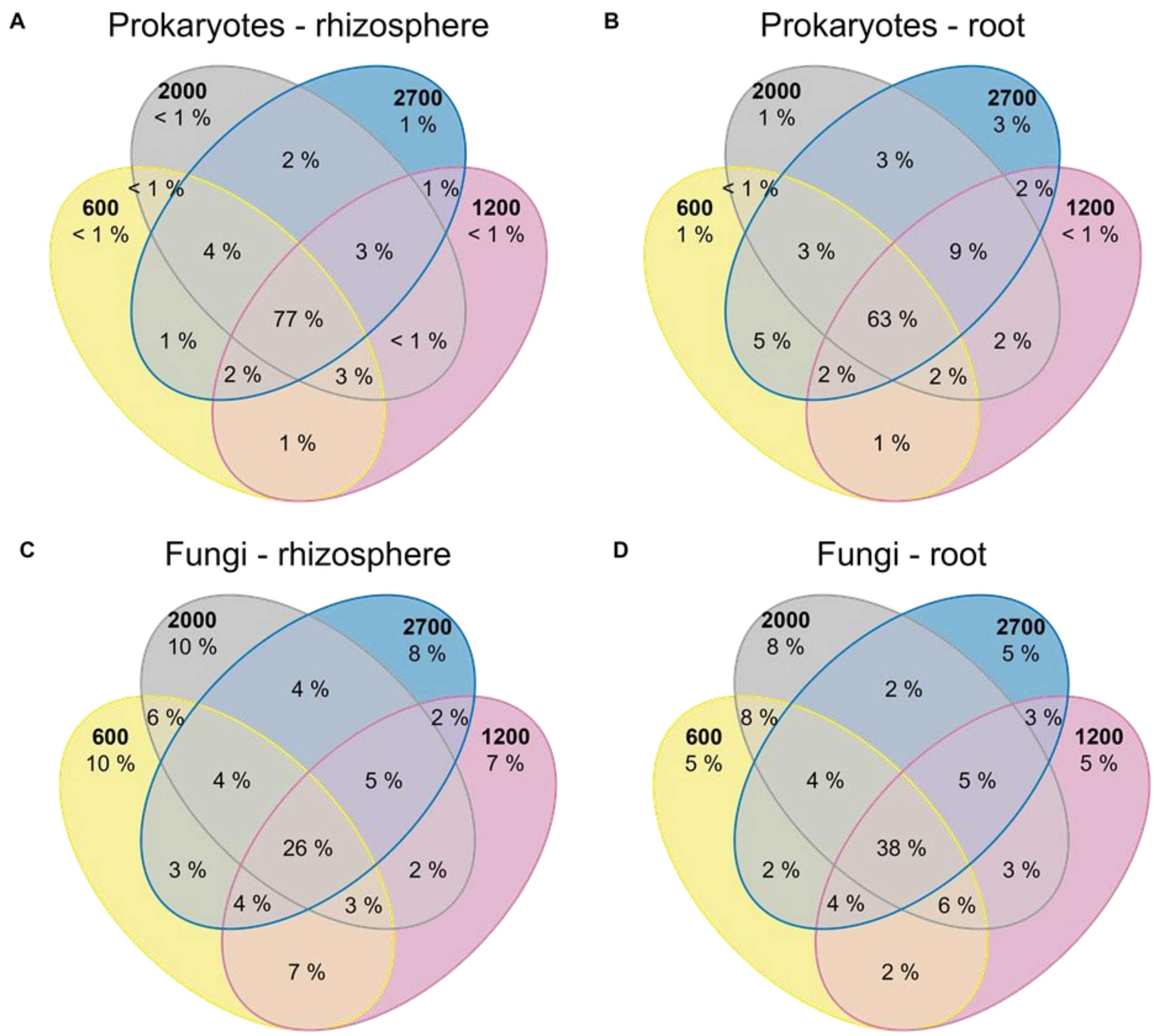


**Fig. S3** Venn diagram showing the overlap in (A) prokaryote ASVs in the rhizosphere, (B) prokaryote ASVs associated to the root, (C) fungal ASVs in the rhizosphere, and (D) fungal ASVs associated to the root between the four levels of mean annual precipitation at locations: 600 mm (yellow), 1200 (pink), 2000 mm (grey), and 2700 mm (blue). Percentages indicate the number of ASVs in each compartment weighted by the ASVs relative abundance (n = 94 per rhizosphere and root compartment).

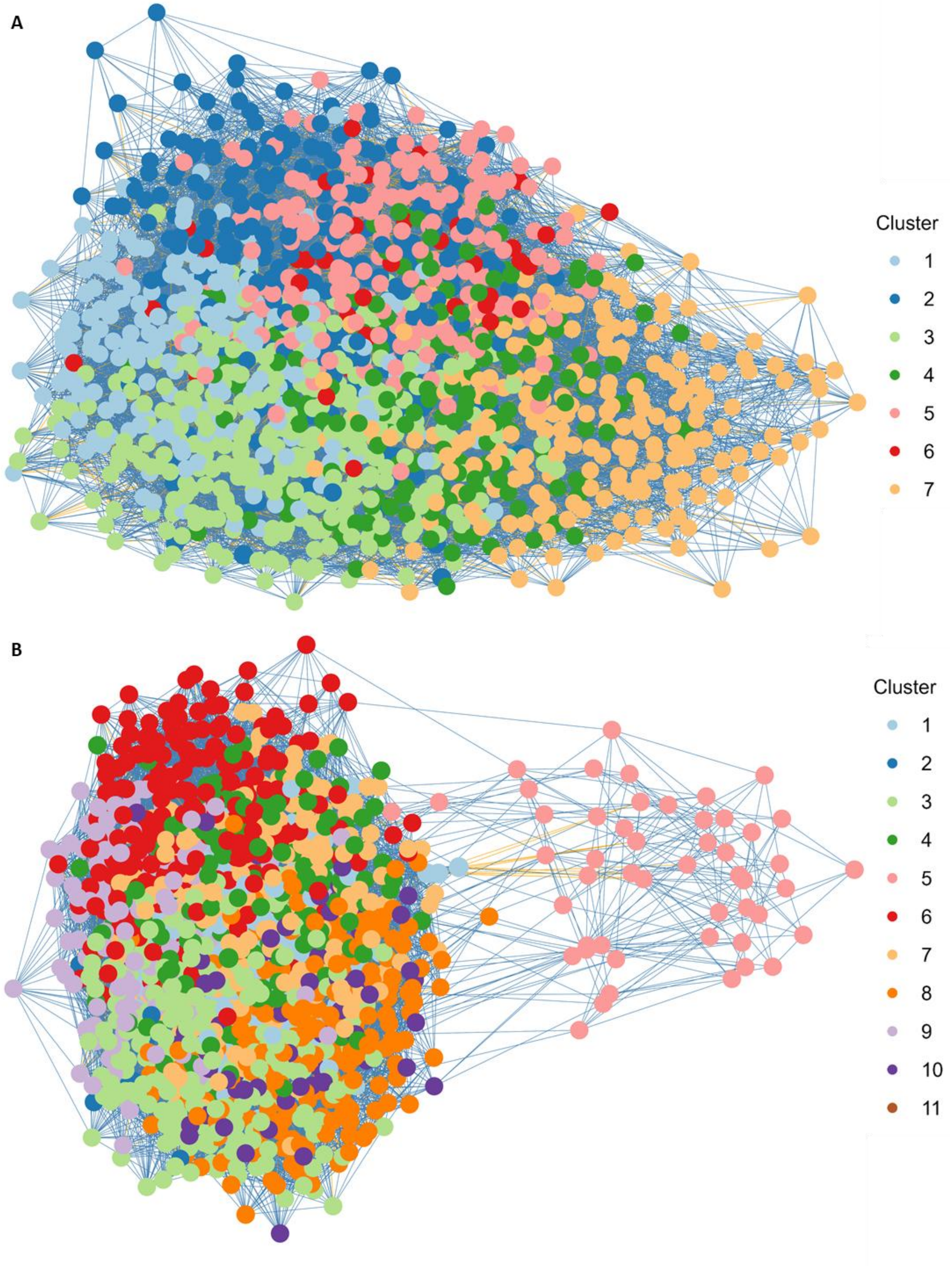


**Fig. S4** *Festuca rubra* (A) rhizosphere and (B) root-associated microbial co-occurrence networks (prokaryotes and fungi together). Each node presents an ASV. Different colours represent different clusters (n = 94).

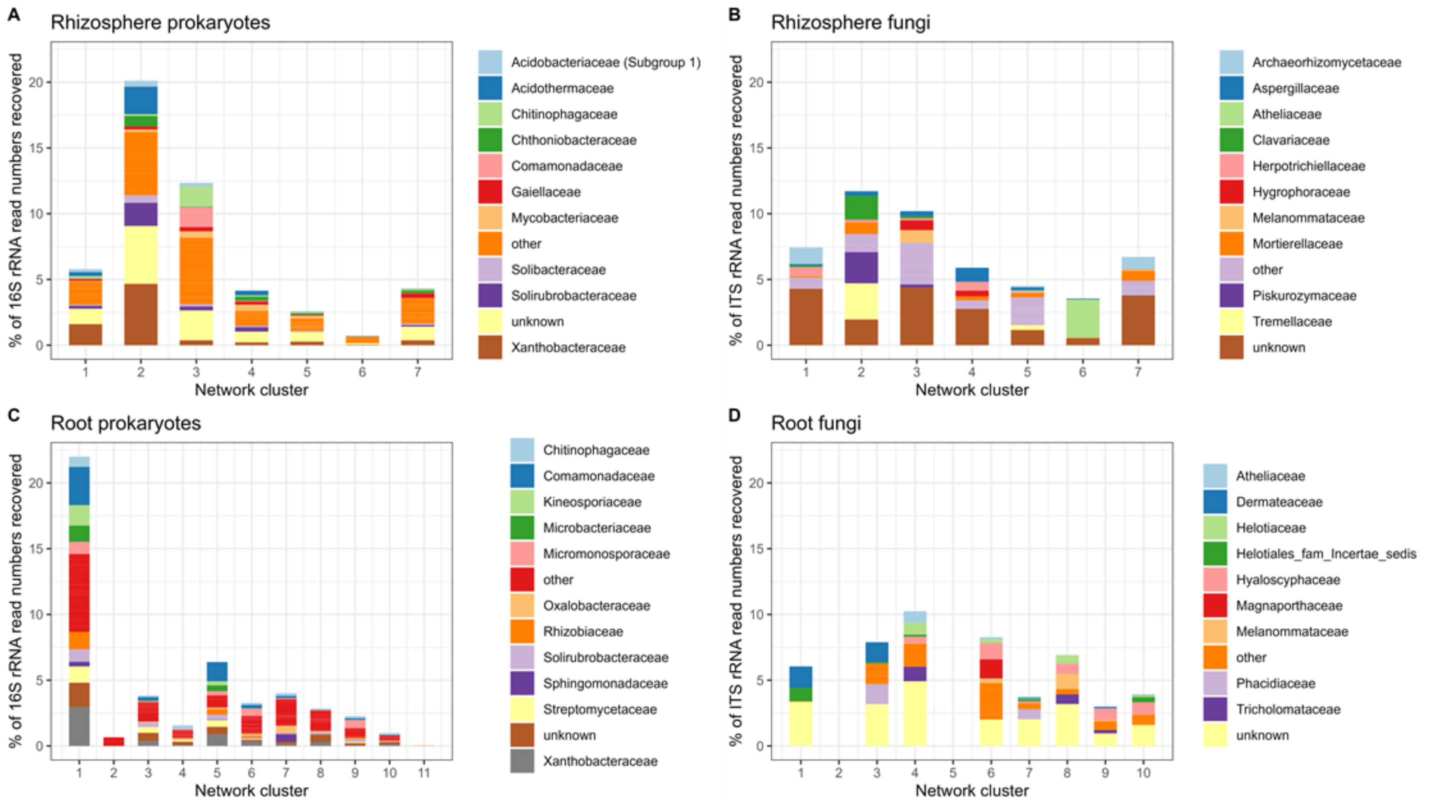


**Fig. S5** Average percentage of 16S and ITS read numbers recovered per family level in prokaryote clusters in *Festuca rubra* (A) rhizosphere soil and (B) roots, and fungal clusters in *F. rubra* (C) rhizosphere soil and (D) roots. Microbial clusters were obtained from co-occurrence networks (Fig 3; S5). For 16S, families < 2% relative abundances are grouped in 'other', for ITS, families <1.5% relative abundances are grouped in 'other'.

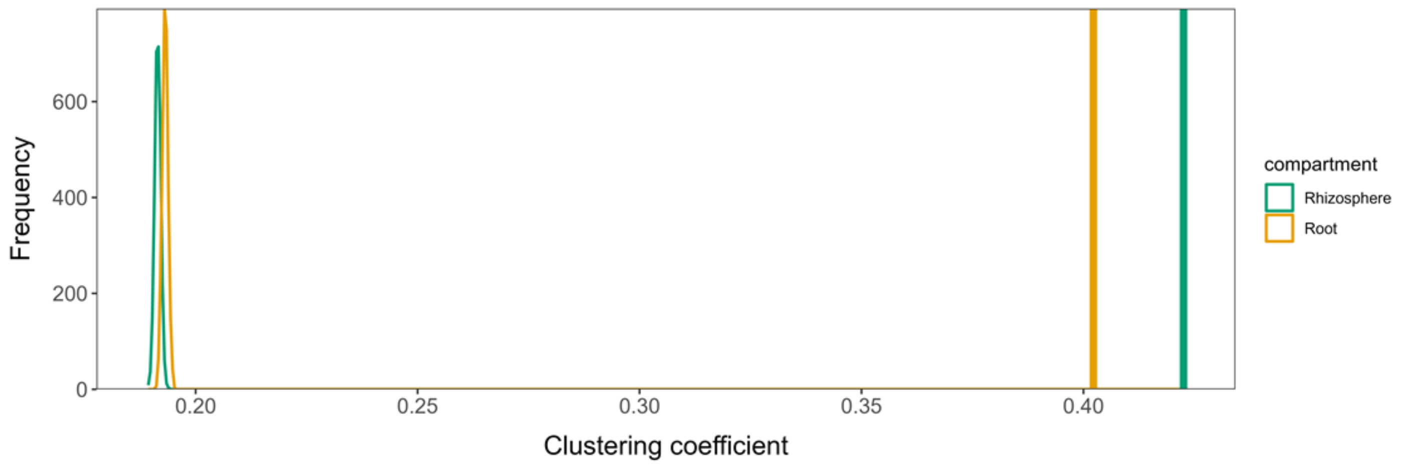


**Fig. S6** Frequency distribution of clustering coefficients (modularity) in random networks of co-occurrence networks from *Festuca rubra* rhizosphere (green) and root (yellow) microbiota. Vertical lines indicate the clustering coefficients of the observed microbial networks. Random networks were created by rewiring the edges of the observed networks while preserving the observed networks degree distribution (1000 iterations). The clustering coefficients of the observed networks did not overlap with the clustering coefficients found for the randomly rewired networks.

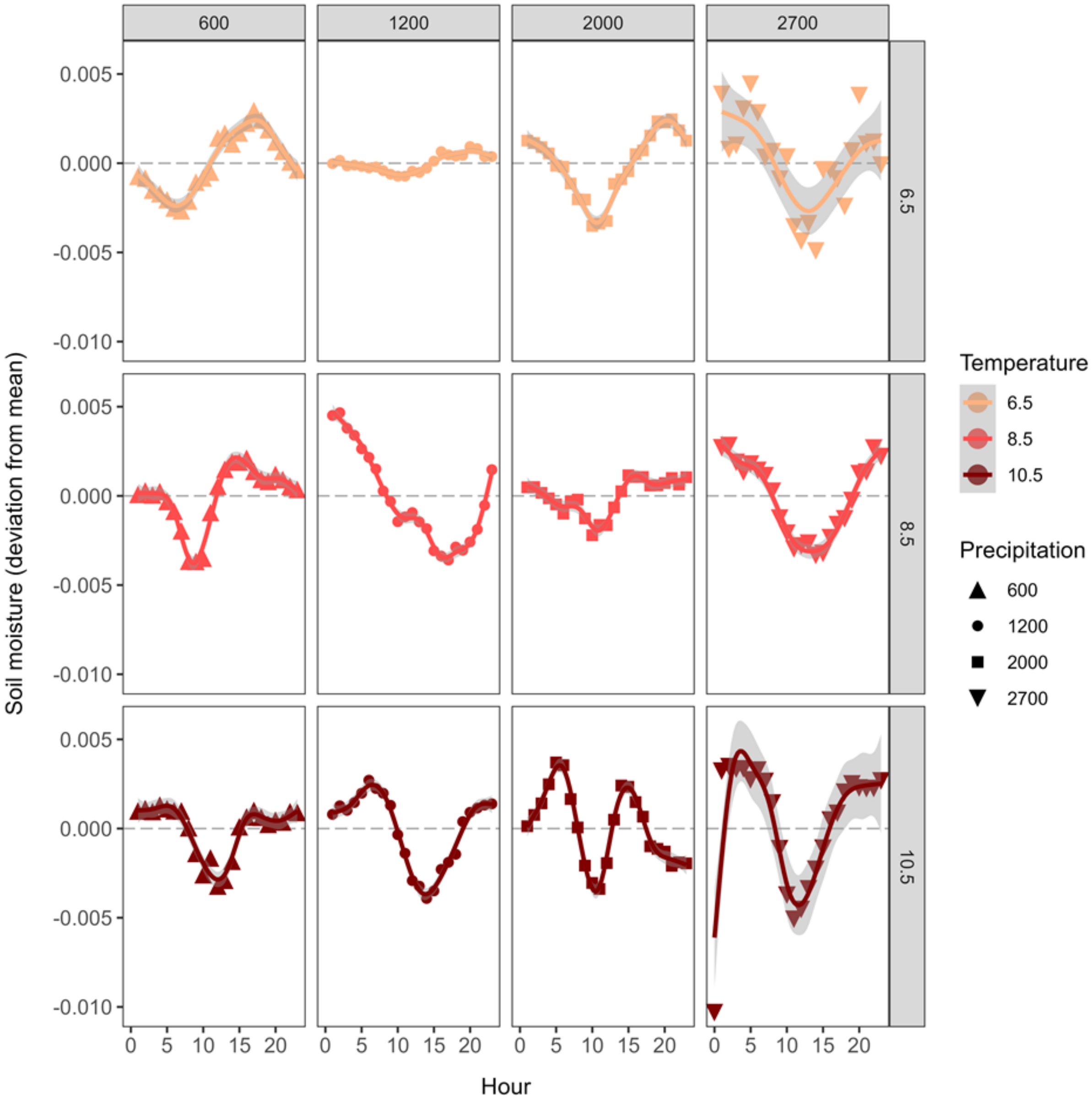


**Fig. S7** Average soil moisture fluctuations during the day, offset to the mean soil moisture at each location. Colours indicate the temperature gradient (average summer temperature in degrees Celsius) and shapes the precipitation gradient (average annual precipitation in mm).

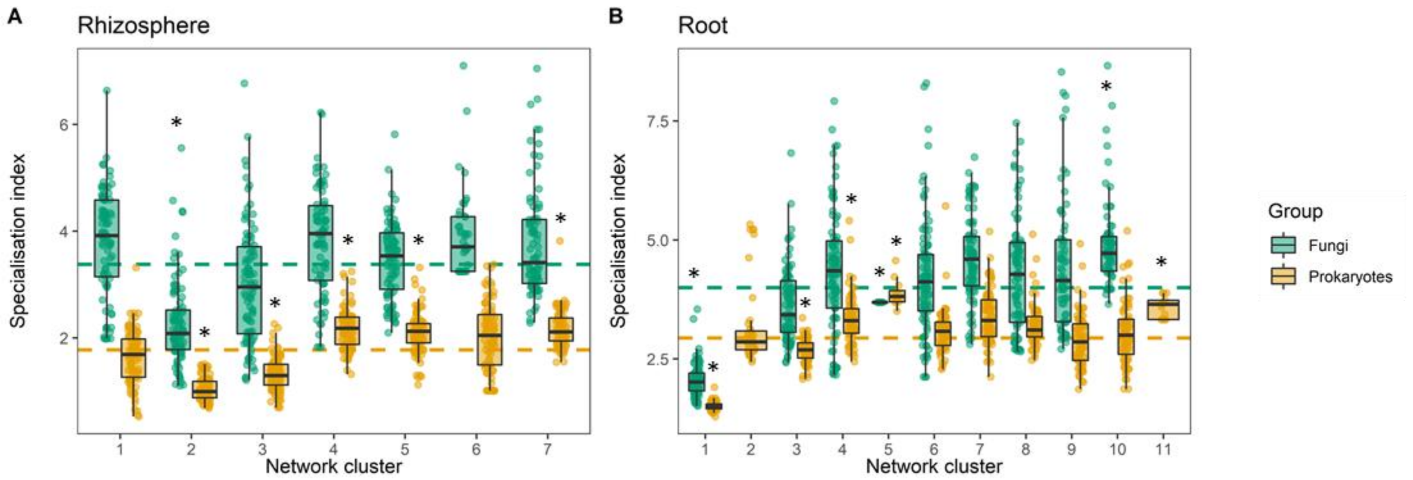


**Fig. S8** Relative habitat specialisation (specialisation index; SI) of (A) rhizosphere co-occurrence networks clusters and (B) root-associated co-occurrence network clusters for fungi (green) and prokaryotes (yellow). Dashed lines indicate the community-wide mean SI of the prokaryote and fungal community, respectively. Relative habitat specialist network clusters occur above the community-wide mean SI without overlap of the minimum distribution (25th percentile - 1.5 * interquartile rang; lower whisker). Relative habitat generalists network clusters occur below the community-wide mean SI without overlap of the maximum distribution (75th percentile + 1.5 * interquartile range; upper whisker). Relative habitat specialist and generalist clusters are indicated by an asterisk (n = 94 samples per cluster).

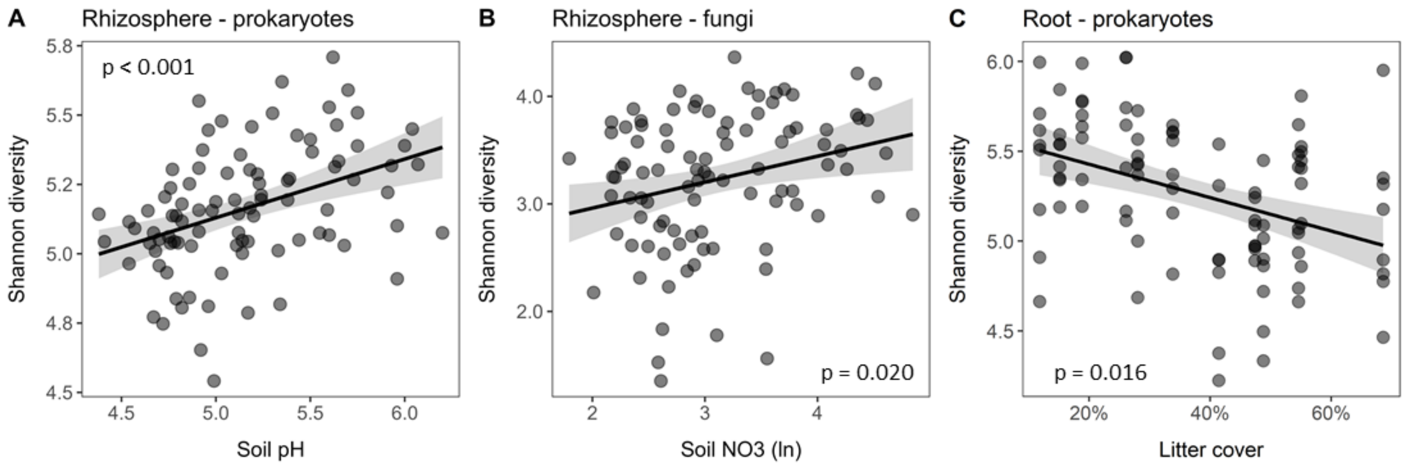


**Fig. S9** Significant relations between Shannon diversity of (A) rhizosphere prokaryotes, (B) rhizosphere fungi, and (C) root-associated prokaryotes to soil properties. Solid lines indicate the mean relation with in grey the 95% confidence interval. Statistical results of the pathways from structural equation models are shown (*n* = 94 *F. rubra* individuals distributed over 12 locations).

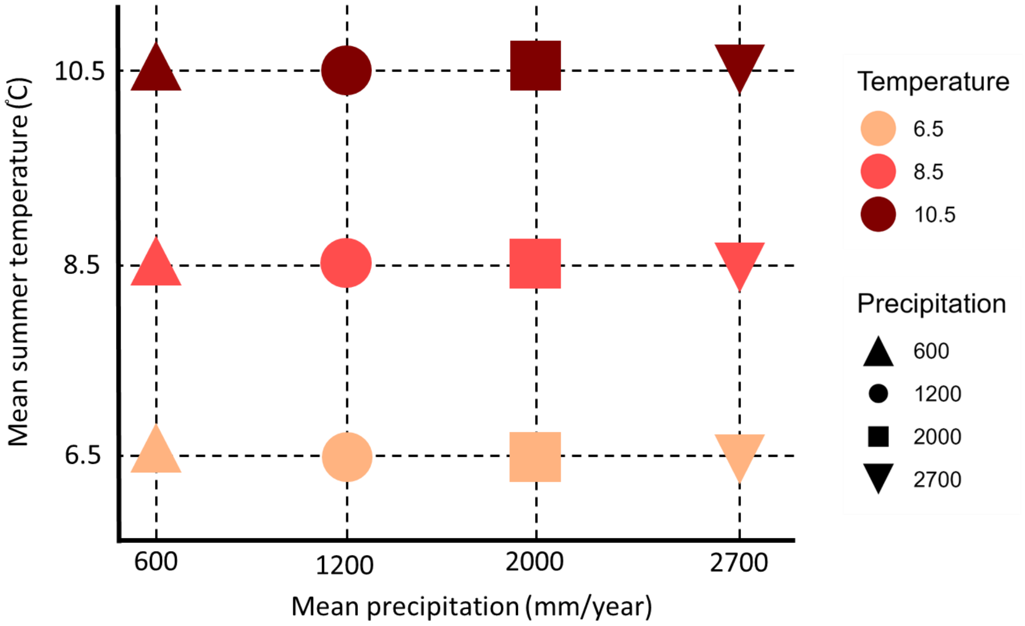


**Fig. S10** Design of the sampled climate grid with three levels of mean summer temperature crossed with four levels of mean annual precipitation. At each location, eight *Festuca rubra* plants were sampled.

### Supplementary tables

**Table S1** Taxonomic composition and fungal traits in microbial network clusters of *Festuca rubra* rhizosphere soil.

| Cluster | Dominant phyla (>10%) | Dominant classes (>10%) | Dominant orders (>5%) | Dominant families (>5%) | Fungal trophic modes (>3%) | Putative function cluster |
|---|---|---|---|---|---|---|
| **1** | ***Ascomycota*** *(44.5%)*<br>*Proteobacteria (21.3%)*<br>***Basidiomycota*** *(10.2%)* | *Alphaproteobacteria (15.9%)*<br>***Leotiomycetes*** *(13.4%)*<br>***Eurotiomycetes*** *(13.1%)*<br>***Agaricomycetes*** *(10.2%)* | *Rhizobiales (12.9%)*<br>***Chaetothyriales*** *(12.3%)*<br>***Helotiales*** *(11.6%)*<br>***Archaeorhizomycetales*** *(9.6%)*<br>*Acidobacteriales (5.2%)* | *Xanthomonadaceae (12.1%)*<br>***Archaeorhizomycetaceae*** *(9.6%)*<br>***Herpotrichiellaceae*** *(5.0%)* | Pathotroph-Saprotroph (5.7%)<br>Saprotroph-Symbiotroph (6.6%) | Widespread, ubiquitous microbiota. Plant pathogens[1], organic matter degraders[2–4], N-fixation and/or phosphate solubilisation[5], acidophilic or acidotolerant nitrite users[6,7]. |
| **2** | *Proteobacteria (28.4%)*<br>***Basidiomycota*** *(22.3%)*<br>*Actinobacteriota (15.5%)* | *Alphaproteobacteria (25.6%)*<br>***Tremellomycetes*** *(16.1%)* | *Rhizobiales (20.1%)*<br>***Tremellales*** *(8.6%)*<br>***Filobasidiales*** *(7.5%)*<br>*Frankiales (6.7%)*<br>***Agaricales*** *(5.7%)*<br>*Solirubrobacterales (5.5%)* | *Xanthomonadaceae (14.7%)*<br>***Tremellaceae*** *(8.6%)*<br>***Piskurozymaceae*** *(7.5%)*<br>*Acidothermaceae (6.6%)*<br>***Clavariaceae*** *(5.7%)*<br>*Solirubrobacteraceae (5.5%)* | Saprotroph (10.3%)<br>Saprotroph-Symbiotroph (9.0%) | Widespread, ubiquitous and diverse microbiota. Slow-growing and depending on recalcitrant organic compounds[8]. Plant pathogens[1], organic matter degraders[9,10], yeasts[11], symbiotrophs[10], N-fixation and/or phosphate solubilisation[5], slow-growing N-fixation[12]. |
| **3** | ***Ascomycota*** *(30.2%)*<br>*Proteobacteria (20.8%)*<br>*Actinobacteriota (16.4%)* | *Gammaproteobacteria (11.8%)* | *Burkholderiales (9.5%)*<br>***Pleosporales*** *(8.3%)*<br>*Chitinophagales (6.9%)* | *Chitinophagaceae (6.9%)*<br>*Comamonadaceae (6.7%)* | Saprotroph (11.4%)<br>Pathotroph-Saprotroph-Symbiotroph (4.2%)<br>Saprotroph-Symbiotroph (4.2%) | Recalcitrant organic matter degradation (a.o. protein/chitin)[13–15]. Heterotrophic ammonia oxidation and denitrification[16]. Aerobic to facultative anaerobic[13,14]. |
| **4** | ***Ascomycota*** *(44.1%)*<br>*Actinobacteriota (22.1%)* | ***Eurotiomycetes*** *(33.9%)*<br>*Actinobacteria (13.1%)* | ***Chaetothyriales*** *(23.3%)*<br>***Eurotiales*** *(10.6%)*<br>*Solirubrobacterales (5.7%)*<br>*Corynebacteriales (5.6%)*<br>*Frankiales (5.4%)* | ***Aspergillaceae*** *(10.6%)*<br>***Herpotrichiellaceae*** *(6.8%)* | Saprotroph (14.9%)<br>Saprotroph-Symbiotroph (7.3%)<br>Pathotroph-Saprotroph (6.8%) | Organic matter degradation and N-fixation. Slow-growing[8,12]. |

Fungi presented in bold.

**Table S1** continued

| Cluster | Dominant phyla (>10%) | Dominant classes (> 10%) | Dominant orders (>5%) | Dominant families (>5%) | Fungal trophic mode (>2%) | |
|---|---|---|---|---|---|---|
| **5** | ***Ascomycota*** *(30.3%)* ***Basidiomycota*** *(25.3%)* *Proteobacteria (13.8%)* | ***Tremellomycetes*** *(13.2%)* ***Agaricomycetes*** *(10.6%)* | ***Tremellales*** *(8.2%)* ***Sordariales*** *(6.1%)* ***Agaricales*** *(5.9%)* ***Helotiales*** *(5.8%)* *Rhizobiales (5.1%)* ***Trichosporonales*** *(5.1%)* *Burkholderiales (5.0%)* | ***Tremellaceae*** *(5.1%)* ***Trichosporonaceae*** *(5.1%)* | Saprotroph (18.0%) Pathotroph-Saprotroph (5.4%) Pathotroph-Symbiotroph (5.1%) Saprotroph-Symbiotroph (4.3%) Pathotroph-Saprotroph-Symbiotroph (3.8%) | Organic matter degradation[17,18]. Yeasts[11], N-fixation and/or phosphate solubilisation [5]. |
| **6** | ***Basidiomycota*** *(68.8%)* ***Ascomycota*** *(13.6%)* | ***Agaricomycetes*** *(68.8%)* | ***Atheliales*** *(68.5%)* ***Helotiales*** *(5.7%)* | ***Atheliaceae*** *(68.5%)* | Pathotroph-Saprotroph-Symbiotroph (68.5%) | Corticoid fungi, organic matter degradation[18,19]. |
| **7** | ***Ascomycota*** *(37.7%)* *Proteobacteria (18.4%)* | *Alphaproteobacteria (11.8%)* | *Rhizobiales (9.5%)* ***Archaeorhizomycetales*** *(8.3%)* ***Helotiales*** *(8.3%)* ***Mortierellales*** *(6.8%)* *Burkholderiales (5.5%)* | ***Archaeorhizomycetaceae*** *(8.3%)* ***Mortierellaceae*** *(6.8%)* | Saprotroph-Symbiotroph (9.7%) Saprotroph (3.6%) | Organic matter degradation[2–4,18]. Require low temperature[20], N-fixation and/or phosphate solubilisation[5]. |

Fungi presented in bold**.**

**References to Table S1**


1. Ryan, R. P. *et al.* Pathogenomics of Xanthomonas: Understanding bacterium-plant interactions. *Nature Reviews Microbiology* vol. 9 344–355 Preprint at https://doi.org/10.1038/nrmicro2558 (2011).

2. Rosling et al. (2011) Science.

3. Thitla, T. *et al.* Exploring diversity rock-inhabiting fungi from northern Thailand: a new genus and three new species belonged to the family Herpotrichiellaceae. *Front Cell Infect Microbiol* **13**, (2023).

4. Fitzpatrick, H. M. *The lower fungi Phycomycetes*. (McGraw-Hill, 1930).

5. Chen, J., Xu, D., Chao, L., Liu, H. & Bao, Y. Microbial assemblages associated with the rhizosphere and endosphere of an herbage, Leymus chinensis. *Microb Biotechnol* **13**, 1390–1402 (2020).

6. Dedysh, S. N. & Oren, A. Acidobacteriales. in *Bergey's Manual of Systematics of Archaea and Bacteria* (ed. Kämpfer, P.) 1–2 (Wiley, 2020). doi:10.1002/9781118960608.obm00001.pub2.

7. Kielak, A. M., Barreto, C. C., Kowalchuk, G. A., van Veen, J. A. & Kuramae, E. E. The ecology of Acidobacteria: Moving beyond genes and genomes. *Frontiers in Microbiology* vol. 7 Preprint at https://doi.org/10.3389/fmicb.2016.00744 (2016).

8. Seki, T., Matsumoto, A., Ōmura, S. & Takahashi, Y. Distribution and isolation of strains belonging to the order Solirubrobacterales. *J Antibiot (Tokyo)* **68**, 763–766 (2015).

9. Berry, A. M., Barabote, R. D. & Normand, P. The Family Acidothermaceae. in *The Prokaryotes* 13–19 (Springer Berlin Heidelberg, 2014). doi:10.1007/978-3-642-30138-4_199.

10. Birkebak, J. M., Mayor, J. R., Ryberg, K. M. & Matheny, P. B. A systematic, morphological and ecological overview of the Clavariaceae (Agaricales). *Mycologia* **105**, 896–911 (2013).

11. Liu, X.-Z. *et al.* Towards an integrated phylogenetic classification of the *Tremellomycetes*. *Stud Mycol* **81**, 85–147 (2015).

12. Normand, P. & Fernandez, M. P. *<scp>F</scp> rankiales* . in *Bergey's Manual of Systematics of Archaea and Bacteria* (ed. Trujillo, M. E.) 1–3 (Wiley, 2020). doi:10.1002/9781118960608.obm00010.pub2.

13. Rosenberg, E. The Family Chitinophagaceae. in *The Prokaryotes* 493–495 (Springer Berlin Heidelberg, 2014). doi:10.1007/978-3-642-38954-2_137.

14. Willems, A. The Family Comamonadaceae. in *The Prokaryotes* 777–851 (Springer Berlin Heidelberg, 2014). doi:10.1007/978-3-642-30197-1_238.

15. Zhang, Y. *et al.* Multi-locus phylogeny of Pleosporales: a taxonomic, ecological and evolutionary re-evaluation. *Stud Mycol* **64**, 85–102 (2009).

16. Wang, Z. *et al.* Impacts of nitrogen-containing coagulants on the nitritation/denitrification of anaerobic digester centrate. *Environ Sci (Camb)* **6**, 3451–3459 (2020).

17. Kendrick, B. *The Fifth Kingdom*. (2000).

18. Watkinson, S. C., Boddy, L. & Money, N. *The Fungi*. (2015).

19. Sulistyo, B. P., Larsson, K.-H., Haelewaters, D. & Ryberg, M. Multigene phylogeny and taxonomic revision of Atheliales s.l.: Reinstatement of three families and one new family, Lobuliciaceae fam. nov. *Fungal Biol* **125**, 239–255 (2021).

20. Alexopoulos, C. J., Mims, C. W. & Blackwell, M. M. *Introductory Mycology*. (The University of California, 1996).

**Table S2** Taxonomic composition and fungal traits in microbial network clusters of *Festuca rubra* roots (endosphere and endoplane)

| Cluster | Dominant phyla (>10%) | Dominant classes (>10%) | Dominant orders (>5%) | Dominant families (>5%) | Fungal trophic mode (>3%) | Putative function cluster |
|---|---|---|---|---|---|---|
| **1** | *Proteobacteria (34.6%)*<br>*Actinobacteriota (35.1%)*<br>***Ascomycota** (21.6%)* | *Actinobacteria (29.8%)*<br>*Alphaproteobacteria (18.6%)*<br>***Leotiomycetes** (16.8%)*<br>*Gammaproteobacteria (16.0%)* | ***Helotiales** (16.8%)*<br>*Rhizobiales (16.1%)*<br>*Burkholderiales (11.8%)*<br>*Kineosporiales (5.6%)* | *Xanthomonadaceae (10.6%)*<br>*Comamonadaceae (10.4%)*<br>***Dermateaceae** (5.8%)*<br>*Kineosporiaceae (5.6%)* | Pathotroph-saprotroph (5.9%) | Mostly plant pathogens[1–3] and disease suppression[4]. Some aerobic organic matter degraders[2,5], N-fixation and anaerobic denitrification[2,6]. |
| **2** | *Firmicutes (99.6%)* | *Bacilli (99.6%)* | *Bacillales (92.1%)*<br>*Paenibacillales (7.5%)* | *Planococcaceae (47.6%)*<br>*Bacillaceae (44.5%)*<br>*Paenibacillaceae (7.5%)* | NA | Plant growth promotion[7–9]. |
| **3** | ***Ascomycota** (50.5%)*<br>*Actinobacteriota (14.4%)*<br>*Proteobacteria (14.0%)* | ***Leotiomycetes** (38.8%)* | ***Helotiales** (25.9%)*<br>***Phacidiales** (12.9%)*<br>*Rhizobiales (6.5%)* | ***Dermateaceae** (13.0%)*<br>***Phacidiaceae** (12.9%)* | Pathotroph-saprotroph (14.5%)<br>Pathotroph-Saprotroph-symbiotroph (3.2%) | Mostly plant pathogens[3,10]. Some organic matter degraders[5], N-fixation and/or phosphate solubilisation[6]. |
| **4** | ***Ascomycota** (50.2%)*<br>***Basidiomycota** (32.2%)* | ***Agaricomycetes** (32.2%)*<br>***Leotiomycetes** (31.2%)* | ***Helotiales** (29.7%)*<br>***Agaricales** (22.6%)*<br>***Chaetothyriales** (8.9%)*<br>***Atheliales** (7.3%)* | ***Tricholomataceae** (9.3%)*<br>***Helotiaceae** (7.8%)* | Pathotroph-Saprotroph-Symbiotroph (20.5%)<br>Pathotroph-Symbiotroph (9.3%)<br>Pathotroph-Saprotroph (4.7%)<br>Saprotroph-Symbiotroph (3.9%) | Related to woody plant species. Corticoid fungi, organic matter degraders[5,11,12], (ecto)mycorrhizae[13,14]. |
| **5** | *Proteobacteria (54.2%)*<br>*Actinobacteriota (38.2%)* | *Actinobacteria (32.7%)*<br>*Gammaproteobacteria (30.2%)*<br>*Alphaproteobacteria (24.0%)* | *Burkholderiales (27.7%)*<br>*Rhizobiales (21.9%)*<br>*Streptomycetales (8.2%)*<br>*Micrococcales (7.0%)*<br>*Solirubrobacterales (5.5%)* | *Comamonadaceae (22.9%)*<br>*Xanthomonadaceae (13.9%)*<br>*Streptomycetaceae (8.1%)*<br>*Microbacteriaceae (7.0%)*<br>*Rhizobiaceae (6.8%)*<br>*Solirubrobacteraceae (5.5%)* | NA | Mostly plant pathogens[1,2,15,16] and disease suppression[17–19]. Some complex organic matter degradation[2,20], N-fixation and anaerobic denitrification[2,6,16]. |

Fungi presented in bold.

**Table S2** continued

| Cluster | Dominant phyla (>10%) | Dominant classes (>10%) | Dominant orders (>5%) | Dominant families (>5%) | Fungal trophic mode (>3%) | Putative function cluster |
|---|---|---|---|---|---|---|
| **6** | ***Ascomycota*** *(57.2%)*<br>*Proteobacteria (11.0%)*<br>*Actinobacteriota (10.2%)*<br>***Basidiomycota*** *(10.0%)* | ***Leotiomycetes*** *(25.1%)*<br>***Sordariomycetes*** *(16.7%)*<br>***Dothideomycetes*** *(11.8%)*<br>***Agaricomycetes*** *(10.0%)* | ***Helotiales*** *(25.0%)*<br>***Magnaporthales*** *(12.6%)*<br>***Pleosporales*** *(11.8%)* | ***Magnaporthaceae*** *(12.6%)*<br>***Hyaloscyphaceae*** *(10.8%)*<br>***Pleosporaceae*** *(5.4%)* | Saprotroph (6.2%)<br>Pathotroph-Saprotroph (5.5%)<br>Pathotroph (5.3%) | Plant pathogens[21,22] and organic matter degraders[5,22,23]. |
| **7** | ***Ascomycota*** *(43.0%)*<br>*Proteobacteria (27.5%)* | ***Leotiomycetes*** *(27.1%)*<br>*Alphaproteobacteria (14.6%)*<br>*Gammaproteobacteria (12.8%)* | ***Helotiales*** *(15.8%)*<br>*Burkholderiales (9.8%)*<br>***Phacidiales*** *(9.7%)*<br>*Polyangiales (7.7%)*<br>*Spingomonadales (7.5%)*<br>*Rhizobiales (6.1%)* | ***Phacidiaceae*** *(9.7%)*<br>*Polyangiaceae (7.7%)*<br>*Sphingomonadaceae (7.5%)*<br>*Oxalobacteraceae (7.0%)* | None | Ubiquitous and diverse microbiota. Plant pathogens[10,24,25], organic matter degraders[5,24–26], growth promotion[24,25], N-fixation and/or phosphate solubilisation[6,25]. |
| **8** | ***Ascomycota*** *(49.2%)*<br>***Basidiomycota*** *(21.7%)*<br>*Proteobacteria (11.1%)*<br>*Actinobacteriota (10.1%)* | ***Agaricomycetes*** *(21.7%)*<br>***Leotiomycetes*** *(21.6%)*<br>***Dothideomycetes*** *(12.3%)* | ***Helotiales*** *(21.4%)*<br>***Pleosporales*** *(11.8%)*<br>***Agaricales*** *(8.4%)*<br>***Auriculariales*** *(5.9%)* | ***Melanommataceae*** *(11.8%)*<br>***Hyaloscyphaceae*** *(7.8%)*<br>***Tricholomataceae*** *(7.5%)*<br>***Helotiaceae*** *(6.9%)* | Saprotroph (8.3%)<br>Pathotroph-Saprotroph (7.6%) | Organic matter degraders[5,23,27], mycorrhizae [13], related to woody vegetation[27]. |
| **9** | ***Ascomycota*** *(28.6%)*<br>*Actinobacteriota (24.4%)*<br>***Basidiomycota*** *(15.4%)*<br>*Proteobacteria (14.0%)*<br>*unknown (13.0%)* | ***Leotiomycetes*** *(21.5%)*<br>*Actinobacteria (20.6%)*<br>***Agaricomycetes*** *(15.4%)*<br>*Gammaproteobacteria (10.1%)* | ***Helotiales*** *(21.5%)*<br>*Micromonosporales (12.0%)*<br>***Trechisporales*** *(10.0%)* | ***Hyaloscyphaceae*** *(17.2%)*<br>*Micromonosporaceae (12.0%)*<br>***Hydnodontaceae*** *(9.5%)* | None | (Complex) organic matter degraders[23,28], true mycelium forming bacteria[28]. |
| **10** | ***Ascomycota*** *(70.6%)* | ***Leotiomycetes*** *(43.5%)*<br>***Sordariomycetes*** *(13.9%)*<br>***Eurotiomycetes*** *(12.5%)* | ***Helotiales*** *(43.5%)*<br>***Cheatothyriales*** *(12.5%)*<br>***Agaricales*** *(5.9%)* | ***Hyaloscyphaceae*** *(19.3%)*<br>***Helotiales fam Incertae sedis*** *(8.0%)*<br>***Strophariaceae*** *(5.9%)* | Pathotroph-Saprotroph (8.8%)<br>Saprotroph (6.9%)<br>Symbiotroph (5.6%) | Organic matter degraders[5,23,29] |
| **11** | *Proteobacteria (100%)* | *Gammaproteobacteria (100%)* | *Burkholderiales (100%)* | *Oxalobacteraceae (100%)* | NA | Diverse. Mostly strictly aerobic, some facultative anaerobic, few strict anaerobic[25]. Saprotrophs, mild plant pathogens and endophytic N-fixation[25]. |

Fungi presented in bold.

#### References to Table S2


1. Ryan, R. P. *et al.* Pathogenomics of Xanthomonas: Understanding bacterium-plant interactions. *Nature Reviews Microbiology* vol. 9 344–355 Preprint at https://doi.org/10.1038/nrmicro2558 (2011).

2. Willems, A. The Family Comamonadaceae. in *The Prokaryotes* 777–851 (Springer Berlin Heidelberg, 2014). doi:10.1007/978-3-642-30197-1_238.

3. Chen, C., Verkley, G. J. M., Sun, G., Groenewald, J. Z. & Crous, P. W. Redefining common endophytes and plant pathogens in Neofabraea, Pezicula, and related genera. *Fungal Biol* **120**, 1291–1322 (2016).

4. Dastogeer, K. M. G., Yasuda, M. & Okazaki, S. Microbiome and pathobiome analyses reveal changes in community structure by foliar pathogen infection in rice. *Front Microbiol* **13**, (2022).

5. Fitzpatrick, H. M. *The lower fungi Phycomycetes*. (McGraw-Hill, 1930).

6. Chen, J., Xu, D., Chao, L., Liu, H. & Bao, Y. Microbial assemblages associated with the rhizosphere and endosphere of an herbage, Leymus chinensis. *Microb Biotechnol* **13**, 1390–1402 (2020).

7. Verma, P. *et al.* Molecular diversity and multifarious plant growth promoting attributes of Bacilli associated with wheat ( *Triticum aestivum* L.) rhizosphere from six diverse agro-ecological zones of India. *J Basic Microbiol* **56**, 44–58 (2016).

8. Mandic-Mulec, I., Stefanic, P. & van Elsas, J. D. Ecology of *Bacillaceae*. *Microbiol Spectr* **3**, (2015).

9. He, C. *et al.* Dual inoculation of dark septate endophytes and *Trichoderma viride* drives plant performance and rhizosphere microbiome adaptations of *Astragalus mongholicus* to drought. *Environ Microbiol* **24**, 324–340 (2022).

10. Crous, P. W., Quaedvlieg, W., Hansen, K., Hawksworth, D. L. & Groenewald, J. Z. Phacidium and Ceuthospora (Phacidiaceae) are congeneric: taxonomic and nomenclatural implications. *IMA Fungus* **5**, 173–193 (2014).

11. Sulistyo, B. P., Larsson, K.-H., Haelewaters, D. & Ryberg, M. Multigene phylogeny and taxonomic revision of Atheliales s.l.: Reinstatement of three families and one new family, Lobuliciaceae fam. nov. *Fungal Biol* **125**, 239–255 (2021).

12. Vrålstad, T., Myhre, E. & Schumacher, T. Molecular diversity and phylogenetic affinities of symbiotic root-associated ascomycetes of the Helotiales in burnt and metal polluted habitats. *New Phytologist* **155**, 131–148 (2002).

13. Campoamor, J. Diversity of Tricholomataceae along a mediterranean altitudinal gradient. *Cryptogam Mycol* **22**, 175–184 (2001).

14. Kernaghan, G. & Patriquin, G. Host Associations Between Fungal Root Endophytes and Boreal Trees. *Microb Ecol* **62**, 460–473 (2011).

15. Evtushenko, L. I. & Takeuchi, M. The Family Microbacteriaceae. in *The Prokaryotes* 1020–1098 (Springer New York, 2006). doi:10.1007/0-387-30743-5_43.

16. Carareto Alves, L. M., de Souza, J. A. M., Varani, A. de M. & Lemos, E. G. de M. The Family Rhizobiaceae. in *The Prokaryotes* 419–437 (Springer Berlin Heidelberg, 2014). doi:10.1007/978-3-642-30197-1_297.

17. Bakker, M. G., Otto-Hanson, L., Lange, A. J., Bradeen, J. M. & Kinkel, L. L. Plant monocultures produce more antagonistic soil Streptomyces communities than high-diversity plant communities. *Soil Biol Biochem* **65**, 304–312 (2013).

18. Cordovez, V. *et al.* Diversity and functions of volatile organic compounds produced by Streptomyces from a disease-suppressive soil. *Front Microbiol* **6**, 1–13 (2015).

19. Kämpfer, P., Glaeser, S. P., Parkes, L., van Keulen, G. & Dyson, P. The Family Streptomycetaceae. in *The Prokaryotes* 889–1010 (Springer Berlin Heidelberg, 2014). doi:10.1007/978-3-642-30138-4_184.

20. Seki, T., Matsumoto, A., Ōmura, S. & Takahashi, Y. Distribution and isolation of strains belonging to the order Solirubrobacterales. *J Antibiot (Tokyo)* **68**, 763–766 (2015).

21. Okagaki, L. H., Sailsbery, J. K., Eyre, A. W. & Dean, R. A. Comparative genome analysis and genome evolution of members of the magnaporthaceae family of fungi. *BMC Genomics* **17**, 135 (2016).

22. Ariyawansa, H. A. *et al.* Towards a natural classification and backbone tree for Pleosporaceae. *Fungal Divers* **71**, 85–139 (2015).

23. Han, J.-G., Hosoya, T., Sung, G.-H. & Shin, H.-D. Phylogenetic reassessment of Hyaloscyphaceae sensu lato (Helotiales, Leotiomycetes) based on multigene analyses. *Fungal Biol* **118**, 150–167 (2014).

24. Glaeser, S. P. & Kämpfer, P. The Family Sphingomonadaceae. in *The Prokaryotes* 641–707 (Springer Berlin Heidelberg, 2014). doi:10.1007/978-3-642-30197-1_302.

25. Baldani, J. I. *et al.* The Family Oxalobacteraceae. in *The Prokaryotes* 919–974 (Springer Berlin Heidelberg, 2014). doi:10.1007/978-3-642-30197-1_291.

26. Garcia, R. & Müller, R. The Family Polyangiaceae. in *The Prokaryotes* 247–279 (Springer Berlin Heidelberg, 2014). doi:10.1007/978-3-642-39044-9_308.

27. Tian, Q. *et al.* Phylogenetic relationships and morphological reappraisal of Melanommataceae (Pleosporales). *Fungal Divers* **74**, 267–324 (2015).

28. Trujillo, M. E., Hong, K. & Genilloud, O. The Family Micromonosporaceae. in *The Prokaryotes* 499–569 (Springer Berlin Heidelberg, 2014). doi:10.1007/978-3-642-30138-4_196.

29. Kirk, P. M., Cannon, P. F., Minter, D. W. & Stalpers, J. A. *Dictionary of the Fungi*. (CAB International, 2008).

**Table S3** Characteristics of microbial network clusters from *Festuca rubra* rhizosphere soil.

| Cluster | Total reads % | Prokaryotes % | Fungi % | Prokaryote unique ASVs | Fungi unique ASVs | SI prokaryote | SI fungi | Relates to |
|---|---|---|---|---|---|---|---|---|
| **1** | 13.2 | 43.8 | 56.2 | 144 | 40 | unspecified | unspecified | Days after defrosting (-); $T_{winter}$ (-); $pH_{soil}$ (-); $PO_4^{3-}{}_{soil}$ (+) |
| **2** | 31.2 | 63.2 | 36.8 | 171 | 48 | generalists | generalists | Days after defrosting (+); Bryophyte cover (-); $pH_{soil}$ (-) |
| **3** | 22.5 | 54.8 | 45.2 | 217 | 63 | generalists | unspecified | Bryophyte cover (-); $pH_{soil}$ (+) |
| **4** | 10.1 | 41.4 | 58.6 | 119 | 39 | specialists | unspecified | Days after defrosting (+); Litter cover (+); $pH_{soil}$ (+) |
| **5** | 7.1 | 36.3 | 63.7 | 97 | 84 | specialists | unspecified | Forb cover (+) |
| **6** | 4.3 | 16.6 | 83.4 | 27 | 10 | unspecified | unspecified | Circadian soil moisture fluctuation (+); $pH_{soil}$ (-); $PO_4^{3-}{}_{soil}$ (-) |
| **7** | 11.0 | 39.0 | 61.0 | 158 | 47 | specialists | unspecified | Soil moisture (+); $pH_{soil}$ (+); $NO_3^-{}_{soil}$ (-) |

The specialisation index (SI) indicates relative habitat generalists and specialists related to variation across all *Festuca rubra* individuals and all locations. Significant relations were obtained from structural equation models.

**Table S4** Characteristics of root-associated microbial network clusters of *Festuca rubra.*

| Cluster | Total reads % | Prokaryotes % | Fungi % | Prokaryote unique ASVs | Fungi unique ASVs | SI prokaryote | SI fungi | Relates to |
|---|---|---|---|---|---|---|---|---|
| **1** | 28.7 | 78.4 | 21.6 | 165 | 10 | generalists | generalists | Precipitation (-) |
| **2** | 0.7 | 100 | 0 | 11 | 0 | unspecified | NA | Days after defrosting (+); Litter cover (-) |
| **3** | 12.0 | 32.7 | 67.3 | 159 | 65 | generalists | unspecified | Days after defrosting (+); $T_{winter}$ (+) |
| **4** | 12.1 | 13.2 | 86.8 | 90 | 48 | specialists | unspecified | Litter cover (+) |
| **5** | 6.5 | 99.9 | 0.1 | 49 | 1 | specialists | generalist | - |
| **6** | 11.8 | 28.4 | 71.6 | 137 | 65 | unspecified | unspecified | $pH_{soil}$ (+) |
| **7** | 7.9 | 51.6 | 48.4 | 147 | 44 | unspecified | unspecified | Days after defrosting (-); Plant diversity (-) |
| **8** | 10.0 | 29.1 | 70.9 | 166 | 47 | unspecified | unspecified | Temperature (-); Bryophyte cover (+); $pH_{soil}$ (-) |
| **9** | 5.4 | 43.2 | 56.8 | 90 | 28 | unspecified | unspecified | Days after defrosting (+); soil moisture (-); $pH_{soil}$ (+) |
| **10** | 5.0 | 20.2 | 79.8 | 44 | 29 | unspecified | specialists | $pH_{soil}$ (-); $NO_3^-{}_{soil}$ (-) |
| **11** | 0.06 | 100 | 0 | 2 | 0 | specialists | NA | - |

The specialisation index (SI) indicates relative habitat generalists and specialists related to variation across all *Festuca rubra* individuals and all locations. Significant relations were obtained from structural equation models.
